# Genome-wide mapping of RPA1 and RAD9 reveals the management of polycistronic transcription, replication initiation, and responses to replication stress in *Leishmania*

**DOI:** 10.1101/2024.11.04.621868

**Authors:** J.A. Black, S. Virgilio, M.S. Bastos, G.L.A. Silva, J.D. Damasceno, C. Lapsley, R. McCulloch, L.R.O. Tosi

## Abstract

When exposed single-stranded DNA accumulates at stalled or collapsed replication forks, the replication stress response is triggered to prevent genome instability. *Leishmania* are parasitic eukaryotes where gene expression is universally polycistronic and whose plastic genomes facilitate rapid adaptations in response to stress, with evidence implicating intrinsic replication stress as a source. Little is known about the *Leishmania* replication stress response. In this study, we reveal the global dynamics of the replication stress response in *L. major* promastigotes by performing ChIP-seq on three key replication stress response proteins, γH2A, RPA1 and RAD9, in the absence and presence of replication stress. We show that common ‘hotspots’ of replication stress correlate with DNA replication initiation and transcription termination in *Leishmania*. When DNA replication is stalled, replication stress response factors accumulate at early S-phase origins, with a signal pattern reminiscent of bidirectional replication fork progression. Under conditions of chronic replication stress, increased accumulation of replication stress response factors emerges at wider sites of transcription initiation, suggesting *Leishmania* may possess compensatory strategies to limit the effects of replication stress and ensure DNA replication can complete under these conditions. In contrast, chronic replication stress enhances RSR factor accumulation at transcription termination sites, highlighting these regions as key replication stress ‘hotspots’ in *Leishmania*. Lastly, variations in RPA dynamics in ATR-deficient cells uncover crucial roles of this protein kinase in managing polycistronic transcription and DNA replication, particularly under replication stress, in *Leishmania*.

**Summary:** Strict controls operate to precisely copy an organism’s DNA. However, cells need ways to rapidly adapt and respond to stimuli. In some cases, these beneficial adaptations come from problems during replication. *Leishmania* parasites cause serious neglected infections in humans and animals across the world’s tropics and sub-tropics. Remarkably, recent evidence suggests that *Leishmania* DNA experiences enhanced stress during replication that can drive its ability to rapidly adapt in response to stress. How L*eishmania* respond to DNA replication stress is still poorly understood. Here, using a genome-wide approach to map the locations of key proteins that manage DNA replication stress and maintain genome integrity, we show ‘hotspots’ of DNA replication stress coincide with start sites of DNA replication and regions of transcription termination.

## Introduction

In most eukaryotes, DNA synthesis begins at the onset of S-phase from multiple DNA replication origins per chromosome licensed during late mitosis and G1 phase[1]. At replication forks (RFs), the replicative helicase moves along the leading DNA strand to unwind the double helix, exposing single-stranded DNA (ssDNA), and the DNA is copied through the actions of DNA polymerases. If these activities become uncoupled, the exposed ssDNA becomes susceptible to nucleolytic attack, base damage, and secondary structure formation that undermine genome stability if unresolved [2]. To preserve ssDNA integrity, single-stranded binding proteins (SSBs) bind to and protect ssDNA, a role primarily fulfilled in eukaryotes by the highly conserved heterotrimeric complex Replication Protein A (RPA), consisting of three subunits [3, 4]. RPA-coated ssDNA then recruits the kinase Ataxia Telangiectasia and Rad3 Related (ATR) and RSR factors including, but not limited to, the heterotrimeric clamp 9-1-1 (RAD9-RAD1-HUS1) and an obligatory interacting partner for ATR (e.g., ATR-interacting partner; ATRIP) protein) [5, 6]. Activated ATR phosphorylates a myriad of downstream factors to halt cell cycle, stabilise RFs and slow origin firing, ultimately limiting the formation of harmful double-strand breaks (DSBs) [6–12]: inhibiting ATR or following prolonged periods of replication stress (RS) induces ssDNA-RPA accumulation, increased origin firing, and a rise in stalled and collapsed RFs [5, 7, 9, 10]. Given the importance of RPA during DNA replication and in the RSR, this complex has been instrumental in mapping the genome-wide distribution of collapsed RFs [7, 13], and wider regions of DNA damage[14]. Mapping the dynamics of yH2A and 9-1-1 during RS has also been used previously to locate damaged RFs and DNA breaks [14, 15].

*Leishmania* are parasitic protozoans that cause the neglected tropical disease (NTD) Leishmaniasis, which is endemic in ~ 90 countries across the tropics and subtropics [16, 17]. Unusually, *Leishmania*, and related kinetoplastids, completely rely on polycistronic transcription of their genome, an arrangement that potentially poses a unique challenge for the management of replication-transcription conflicts [18]. Marker Frequency Analysis followed by next generation sequencing (MFA-seq) revealed a single early-S replication initiation site per chromosome, located within a strand switch region (SSR) rich in acetylated histone H3 (AcH3), Base J, and the kinetochore factor KKT1 [19], suggesting these sites additionally support centromere formation[20]. Moreover, MFA-seq also supports the co-orientation of the leading strand with transcription, potentially enabling *Leishmania* to avoid machinery collisions within the chromosome cores [20]. Mapping origin sites using Single Molecular Analysis of Replicated DNA (SMARD) revealed multiple origins per chromosome, but their locations relative to transcriptional activity are unknown [21] whereas Short Nascent Strand (SNS-seq [22]) suggests that *Leishmania* chromosomes possess hundreds of origins, though these sites overlap with features related to mRNA maturation, leaving their role in replication initiation unclear. To date, no study has fully described the *Leishmania* DNA replication program or its interactions with the parasite’s multigenic transcription landscape. In addition, little is known about how *Leishmania* manage interactions between DNA replication and transcription machineries. Genomic regions at the ends of multigenic units (Strand Switch Regions; SSRs), are used to regulate both replication and transcription activities [19, 22, 23]. Transcription initiation sites are marked by acetylated histone H3 (AcH3;[24–26]) and transcription termination sites by the glycosylated thymidine base J [24, 27–29]. However, little is known about the RSR in *Leishmania*. The *Leishmania* genome encodes a complete 9-1-1 complex, RPA complex, and the ATR kinase, suggesting core aspects of the RSR are retained, with functional studies of these factors revealing roles in cell cycle progression, RS and the response to DNA damage [30, 31]. However, *Leishmania* appears to lack a CHK1 homolog (downstream target of ATR) [32, 33] and additionally harbours divergent 9-1-1[31, 34] and RPA functions, raising questions as to the specific dynamics of their RSR. In fact, the remarkable plasticity endured by the *Leishmania* genome, such as extensive chromosome and gene copy number alterations [35–39], single nucleotide polymorphisms, and the production of extrachromosomal DNA elements in the form of linear or circular molecules of DNA [40–43], suggests otherwise, indicating a less stringent RSR, particularly given subtelomeric duplication may require components of the 9-1-1 complex in *Leishmania* [19].

RPA1 in *Leishmania* is required for telomere protection, the DNA damage response and ssDNA binding [44–48], whereas investigations into RPA2, in *T. brucei* reveal functions associated with both DSBs [49]and the RSR[50] suggesting the RPA complex has broad functions pertaining to genome stability in these parasites. However, given RPA1’s demonstrated capacity to interact and bind ssDNA in *Leishmania*, we reasoned that RPA1, as a proxy for the RPA complex, would enable the global profiling of the interactions between a protein that is located directly at RFs and is functionally linked to the RSR in *Leishmania*. In this study, we performed ChIP-seq on *L. major* promastigote cells with endogenously tagged RPA1, in the absence and presence of acute and chronic conditions of RS induced by Hydroxyurea (HU). To assess genotoxic stress and putative sites of DNA damage, we additionally mapped the locations of RAD9 (9-1-1 complex) and yH2A, a marker of DNA damage [51, 52]. Additionally, to investigate the process under sustained RS, we analysed RPA1 dynamics in an ATR-deficient mutant.

We show that RPA1 accumulates at DNA replication sites during early-S phase, in keeping with previous studies [19, 53]. Altogether, our work highlights key events of replication-transcription conflict or replication initiation found at the boundaries of polycistronic transcription units (PTUs), reflecting the unique organisation of gene expression in this and related kinetoplastids. The relative lack of such interactions within the PTUs suggests undetected mechanisms to minimise conflicts between actively moving DNA replication and transcription machineries. The data may also suggest a means to ensure copying of at least the larger chromosomes during RS conditions. Furthermore, variations in peak sizes and signal distribution in ATR-deficient cells reveals key roles for this kinase in the management of multigenic transcription and genome duplication, exemplified by induction of RS, in *L. major* promastigotes.

## Results

### *Leishmania* RPA1 forms nuclear foci under Replication Stress

To investigate whether RPA1 responds to RS in *L. major*, we performed immunoblotting on total protein extracts from cells exposed to either acute (5 mM; 2 and 8 hrs) or chronic (0.5 mM; 8 and 20 hrs) HU treatment and probed for RPA1 (using *L. major* anti-RPA1) and yH2A (using *T. brucei* anti-yH2A) [51]. Both acute and chronic HU treatments increased the levels of yH2A, relative to controls, indicating enhanced DNA damage due to HU stress. However, the total protein levels of RPA1 remained unchanged compared with controls (**Fig. 1A; SFig. 1A**). We hypothesized that under replication stress, the RPA complex might reorganize into nuclear foci, similar to the behaviour of RPA after exposure to DNA damage in *T. brucei* [49, 50, 54] and *T. cruzi* [55]. To investigate RPA1 foci formation in *L. major* cells, we generated a 3xMyc-RPA1 cell line (referred to as RPA1myc) by fusing three copies of the Myc epitope to the N-terminus of RPA1 using CRISPR/Cas9 genome editing (**Fig. S1B**). PCR confirmed the correct integration of the tag into the locus (**Fig. S1C**), and immunoblotting with anti-Myc (**Fig. S1D; left**) and anti-RPA1 (**Fig. S1D; right**) antibodies verified the expression of the fusion protein at the expected molecular mass (~56 kDa). RPA1myc cells did not show any significant growth or cell cycle defects relative to controls (**Fig. S1E&F; Supplementary Data 3**), indicating expression of the fusion protein does not affect parasite proliferation.

**Figure 1.**
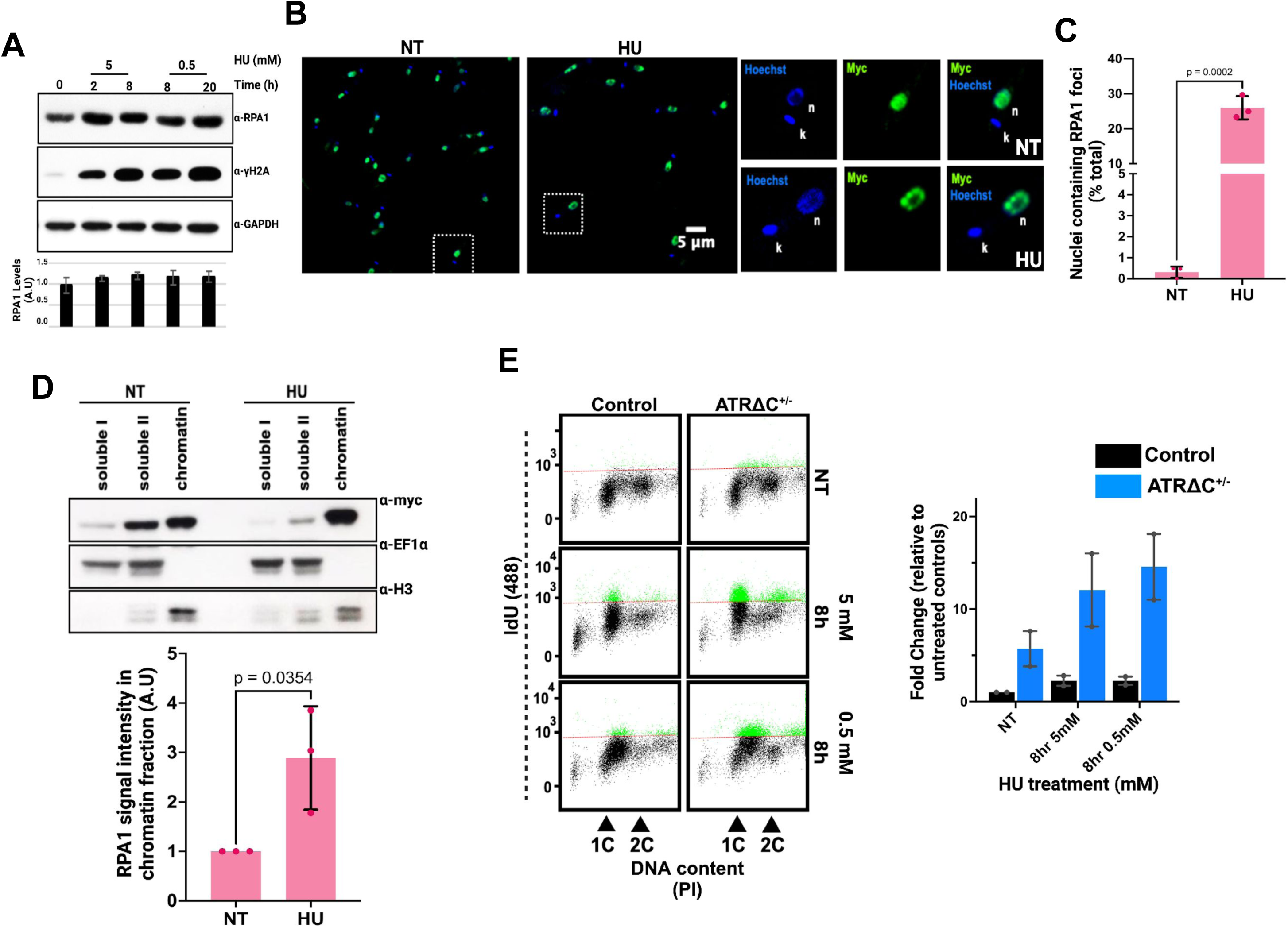
RPA1 forms nuclear foci in the presence of replicative stress. (A) Immunoblot showing RPA1 and yH2A signal after exposure to HU at two concentrations (0.5 mM and 5 mM) at the times shown in the figure. The membrane was probed with anti-LmRPA1 and anti-yH2A antiserum and Anti-GADPH was used as a loading control. Graph below shows the RPA1 signal quantified relative to the loading control (displayed in arbitrary units; A.U). (B) Representative indirect immunofluorescence images of *L. major* parasites expressing RPAmyc in the absence (left) or presence (right) of replication stress induced by the addition of 5 mM Hydroxyurea (HU) for 10 hrs. RPAmyc was probed for using anti-myc antiserum (green) and Hoechst used to visualise the nuclear and kinetoplastid DNA (blue). NT = non treated, dashed white boxes indicate the regions enhanced from the field of view. k = kinetoplast, n = nucleus. Scale bar = 5 μm. Images were captured on a Zeiss Multiphoton confocal microscope. (C) The percentage of RPAmyc cells with nuclear RPA1 foci in the absence and presence of replication stress (5 mM HU for 10 hrs). Error bar = +/− SD, n = 3 independent experiments. (D) Chromatin extraction performed on RPAmyc cells in the presence or absence of 5 mM HU treatment (4 hr exposure). Histone H3 (H3) was used to confirm isolation of the chromatin fraction and EF1alpha as a loading control for all other fractions. Anti-myc antiserum was used to detect RPAmyc. Gel is representative of 3 independent experiments. The intensity of the myc signal from chromatin fraction was quantified in the absence and presence of HU. The graph shows the relative intensity of myc signal in the HU treated cells relative to controls (NT). Signals are shown as arbitrary units (A.U.). Error bars = +/− SD, n = 3 independent experiments. (E) Quantification of ssDNA levels by flow cytometry under native conditions in the absence and presence of RS induced by HU at 8hrs (0.5 mM) and 8 hrs (5 mM). The fold changes of ssDNA intensity relative to unstained negative controls for each sample are shown in the graph (right). Error bars = +/− SEM, n = 2 independent experiments.

In unperturbed RPA1myc cells, indirect immunofluorescence analysis (IFA) revealed a diffuse Myc signal (green) localizing exclusively at the location of the larger DAPI signal (blue) from the cell nucleus (**Fig. 1B; left panel; Fig. S1G and H** show untagged control cells), confirming that RPA1 is a nuclear protein in *L. major*. When parasites were subject to replication stress conditions via treatment with 5 mM HU for 10 hours, we detected distinct nuclear foci of Myc signal in approximately 25% of the cells, compared to less than 1% in untreated cells (**Fig. 1B, right panel; Fig. 1C**). The presence of nuclear RPA foci suggests enhanced recruitment of RPA1 to the chromatin. To assess whether the 3xMycRPA1 protein becomes chromatin-enriched under replication stress, we performed chromatin fractionation with and without HU treatment. Following the acute HU treatment, we observed an ~3-fold increase in Myc signal associated with the chromatin fraction, while the signal in other soluble fractions decreased (**Fig. 1D**; **Fig. S1I**). These findings demonstrate that RPA1 in *L. major* becomes enriched on chromatin and forms nuclear foci in response to HU treatment.

### ATR-deficiency enhances replication stress in *Leishmania*

Accumulation of RPA-coated ssDNA promotes activation of ATR in response to RS[2, 6]. Considering that ATR inhibition leads to excessive origin firing and accumulation of ssDNA in human cells [9], we investigated the dynamics of RPA1 in an ATR-deficient RPA1myc cell line. We opted to truncate one allele of ATR in RPA1myc cells by removing a C-terminal proportion of the ORF encoding the kinase domain using CRISPR/Cas9 editing, generating the cell line 3xMycRPA1^ATRDC+/−^ (referred to hereafter as ATRΔC+/−) (**Fig. S2A**). Truncation of one allele of ATR in the ATRΔC+/− cells was confirmed by PCR (**Fig. S2B)** and Southern blotting (**Fig. S2B**). We found no significant alteration to parasite proliferation in ATR-deficient cells, nor enhanced sensitivity to HU (**Fig. S2D&E**). To determine if ATRΔC+/− cells show other signs of ATR deficiency, we asked if ATRΔC+/− cells accumulate ssDNA by assessing ssDNA levels using flow cytometry (**Fig. 1E**) and IFA (**Fig. S2F&G**) through the detection of incorporated IdU (a thymidine analogue) under native conditions. In the absence of HU-induced replication stress, we detected a ~ 5-fold increase in ssDNA levels in ATRΔC+/− cells compared with control cells (**Fig. 1E**). After HU exposure, the percentage of ssDNA positive cells significantly increased compared with untreated controls at 8 hrs 5 mM HU (acute) and 8 hrs 0.5 mM HU (chronic). A similar trend was observed by IFA (**Fig. S1F&G**) where ssDNA positive cells increased in ATRΔC+/− cells compared with controls after HU exposure. In addition, we detected a significant increase in the fluorescence intensity of the RPAmyc signal in ATRΔC+/− cells, relative to controls, which significantly increased under conditions of RS (**Fig. S2G**). Altogether, these findings suggest that ATR deficiency enhances replication stress in *L. major*, even in the absence of HU.

### Genome-wide mapping of yH2A, RPA1 and RAD9 in unperturbed *L. major* reveals RPA1 at sites of DNA replication initiation

To delinate the RSR in *Leishmania*, we first sought to map the genome-wide distribution of RPA1 in asynchronously dividing promastigotes by ChIP-seq, targeting the 3xMycRPA fusion protein expressed in both RPA1myc and ATRΔC+/− cells using ChIP-grade anti-myc antiserum. To evaluate the implications of the localization, we also investigated the global dynamics of yH2A using anti-yH2A antiserum, aiming to identify sites of pronounced replication stress and/or DNA damage[15]. Additionally, we examined the location of the 9-1-1 complex, which is recruited to damaged RFs and DNA breaks under RS conditions[14, 56]. We also used RAD9 as a proxy for the 9-1-1 complex by performing ChIP-seq in a cell line expressing 12xMyc epitope fused to the C-terminus of RAD9 (RAD9myc[57]). Two independent experiments were performed for each 3xMyc-RPA1-expressing cell line and for the yH2A ChIP-seq, with high reproducibility between replicates (**Fig. S3A-D**). ChIP-sep of RAD9 was performed once, with ChIP-seq of the HUS1 subunit provided in **Supplementary Data 1** as support for the locations of RAD9. Whole chromosome plots are provided in **Supplementary Data 2**.

RPA1 ChIP signals from RPA1myc and ATRΔC+/− cells and RAD9 ChIP signal from RAD9myc cell line were strongly correlated (Pearson coefficient: 0.88 - 0.92; **Fig. S3B**), suggestive of a similar enrichment pattern. To investigate this correlation further, we centered the ChIP signal from RPA1-enriched regions in RPA1myc cells (defined by MACS2) and analyzed the profile of yH2A, RAD9, and RPA1 in ATRΔC+/− cells in relation to these RPA1-bound sites. We used k-means clustering to group regions with similar signal patterns for γH2A, RPA1, and RAD9. Consequently, we identified three distinct signal clusters, encompassing a total of 89 enriched regions (**Fig. 2A**). In cluster 1, which exhibited the strongest and broadest RPA1 enrichment, approximately 97% of the regions overlapped with the previously mapped early-S phase DNA replication initiation sites[19, 48], suggesting that one of the major sites of RPA accumulation are coincident with single DNA replication origins detected by MFA-seq in S-phase in *L. major*. These sites also showed co-enrichment with RAD9, indicating RS and/or DNA damage at DNA replication initiation sites. We found no evidence of yH2A accumulation upstream or downstream of the early-S DNA replication origins (**Fig. 2A**). Cluster 2, which also showed both RPA and RAD9 accumulation, primarily matched transcription termination sites, with approximately 80% of the enriched regions overlapping with sites decorated with BaseJ alone or by both BaseJ and AcH3. SNS-seq predicted DNA replication initiation at transection termination sites that was not detected by MFA-seq[22]; since these sites accumulate both RPA and RAD9, it is more likely that replication stress and not DNA replication initiation is being detected, perhaps suggesting that transcription termination sites are ‘pronounced sites of RS. Only ~ 6% of the enriched regions in cluster 2 overlapped with AcH3 alone (sites associated with transcription initiation only). Lastly, in cluster 3, which exhibited the weakest enrichment signals, approximately 62% of the regions overlapped with various other genomic features including inter-CDS sites and UTRs, indicating wider sites of DNA instability and/or repair which could be associated with trans-splicing activity and/or polyadenylation. These regions also accumulate RNA-DNA hybrids in both *Leishmania*[58] and *T. brucei*[59], and are therefore likely more prone to elevated levels of ssDNA formation.

**Figure 2.**
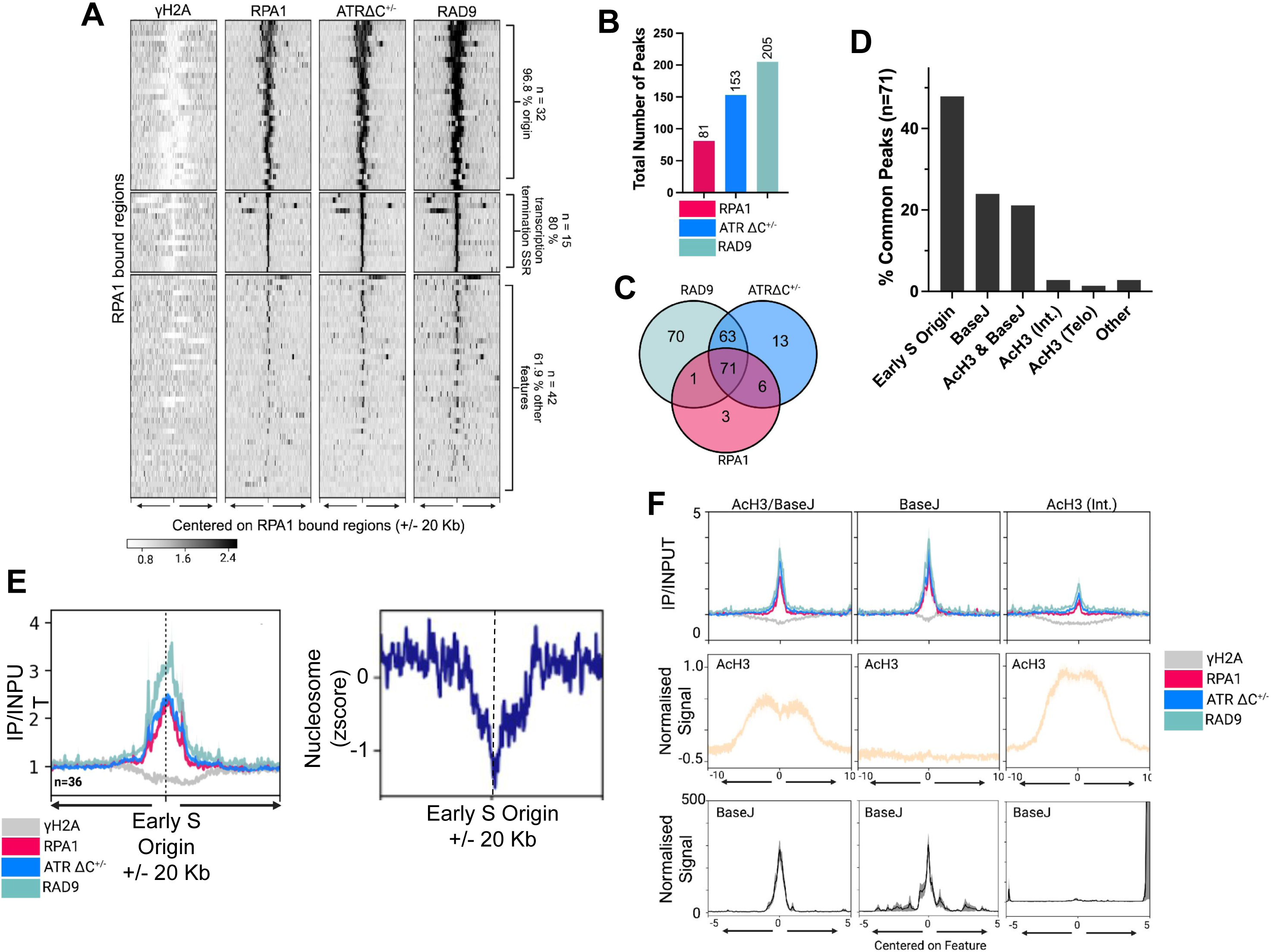
Genome-wide mapping of yH2A, RPA1 and RAD9 dynamics in unperturbed cells. (A) RPA1-bound sites were identified using MACS2 then the signals for yH2A, RPA1, RPA1 (in ATR-deficient cells) and RAD9 were centred on these sites +/− 20 Kb up- and down-stream of the regions. The colour white on the heatmap shows no binding. K-means clustering was used to group sites of similar signal together and Bedtools intersect used to locate the genomic feature associated with these RPA1-bound sites. (B) The total number of peaks for RPA1 and RAD9 identified using MACS2. Peaks were filtered for quality (> 20) and removed if associated with regions of high coverage (‘shade regions’). (C) RPA1, in ATR competent and ATR deficient cells, and RAD9 co-occupy 71 genome regions in unperturbed cells. (D) The genomic features associated with these 71 common peaks are shown as a percentage of the total number of common peaks. (E) Metaplot analysis depicts enriched signal for yH2A, RPA1 and RAD9 centred on early-S phase origins +/− 20 kb. Immunoprecipitated samples were normalised to their corresponding input samples. Metaplot to the right shows the nucleosome content of the region. (F) Metaplot analyses depicting enriched signal for yH2A, RPA1 and RAD9 centred on sites enriched for AcH3 (transcription initiation - chromosome cores), BaseJ (transcription termination) and AcH3/BaseJ (transcription initiation and termination) +/− 5 kb.

MACS2 was then used to call peaks across all samples, after which the peaks were filtered to exclude regions of high coverage (‘shade areas’[22]), resulting in the identification of 81 RPA1 peaks in RPA1myc cells and 153 RPA1 peaks in ATRΔC+/− cells, as well as 205 RAD9 peaks in the RAD9myc cell line (**Fig. 2B&C**). The approximately 1.8-fold increase in RPA1 peaks in ATRΔC+/− cells compared to the RPA1myc cell line aligns with the increased ssDNA signal observed in earlier data (**Fig. 1E and Fig. S2F&G**), indicating heightened RS and/or damage when ATR is deficient. Notably, of the 63 common RPA1 and RAD9 peaks found exclusively in ATRΔC+/− and RAD9myc cells, over 50% (33 peaks) co-localized with SSRs associated with transcription initiation, including 8 peaks located at subtelomeric sites **(Fig. S3E**). This suggests that ATR function is particularly relevant for stabilizing these specific regions. Unique RPA1 peaks were found in RPA1myc (3 peaks) and ATRΔC+/− cells (13 peaks), while unique RAD9 peaks were observed in RAD9myc cells (70 peaks). These unique peaks were associated with various genomic features (**Fig. S3F**), including intergenic regions and gene coding sequences, which we propose reflect spontaneous genome instability events across the genome. Consistent with the data shown in **Fig. 2A**, nearly all the common RPA1 and RAD9 peaks (71) identified in both RPA1myc and ATRΔC+/− cells (**Fig. 2D**) were located at the boundaries of polycistronic transcription units (PTUs). Specifically, 48% of these peaks (34) were associated with early-S origins. Additionally, 49% (32) were found at sites of transcription termination (head-to-tail and convergent SSRs). Only about 4% of the common peaks were located at sites of transcription initiation, either within the interior of chromosomes (divergent SSRs, 2.8%; **Fig. 2D**) or at subtelomeres (AcH3 Telo, 1.4%; **Fig. 2D**)

Metaplot analysis confirmed broad enrichment of each protein at the early-S origins (**Fig. 2E**), with RPA1 signal from RPA1myc and ATRΔC+/− cells largely overlapping. This suggests that ATR deficiency does not affect RPA1 accumulation at early-S DNA replication initiation sites in unperturbed cells. We noted a reduction in yH2A ChIP signal at the center of these sites, and little accumulation 20 kb +/− the regions, likely because these replication initiation sites are nucleosome-depleted **(Fig. S3E**), preventing the generation of yH2A modification. In contrast, metaplot analyses of the other SSRs revealed sharp peaks for RPA and RAD9, but not yH2A, at sites where transcription initiates and/or terminates (AcH3/BaseJ or BaseJ sites) with markedly reduced peak sizes at transcription initiation sites (AcH3). (**Fig. 2F**). No significant accumulation of either RPA or RAD9 was detected across the subtelomeres (**Fig. S3G**). However, a modest increase in RPA1 (across all cell lines) and RAD9 signal was observed at subtelomere-proximal transcription initiation sites (**Fig. S3H**).

Taken together, these data show that in the absence of exogenous RS, RPA1 and RAD9 predominantly accumulate at PTU boundaries, widely at early origins and more focused at transcription termination sites, highlighting these regions as the primary locations of RS and/or DNA damage and repair that are clearly detectable in *Leishmania*. Additionally, whilst ATR deficiency had a minimal impact on asynchronously dividing cells, the marked increase of RPA at transcription initiation sites in ATRΔC+/− cells underscores these regions as damage ‘hotspots’ when ATR function is impaired.

### RPA1 accumulates bidirectionally from early-S origins under acute replication stress in *L. major*

In unstressed *L. major* promastigotes, where most of the population is in G1 phase, approximately half of sites enriched with RPA1 and RAD9 signal localized to the previously mapped single site of DNA replication initiation on each chromosome[19, 48], suggesting much of this observed signalling response is primarily associated with DNA replication activities. To next investigate the dynamics of the RSR, we examined the localisation of yH2A, RPA1 and RAD9 following HU treatment, which inhibits dNTP synthesis and induces RS[60]. We first treated RPA1myc, ATRΔC+/− and RAD9myc cells with 5 mM HU for 2 hrs and 8 hrs (**Fig. 3A**). Under these conditions, the *L. major* cell cycle is stalled during G1/early-S [31]. We detected a strong correlation between replicates (**Fig. S5A**). Like untreated cells, normalized ChIP signals from RPA1myc, ATRΔC+/− and RAD9myc cells after 2 hrs and 8 hrs of HU exposure were highly correlated, indicating a consistent enrichment pattern (Pearson coefficient: 0.69-0.97; **Fig. S5B**). Full chromosome signal plots are shown in **Supplementary Data 3.**

**Figure 3.**
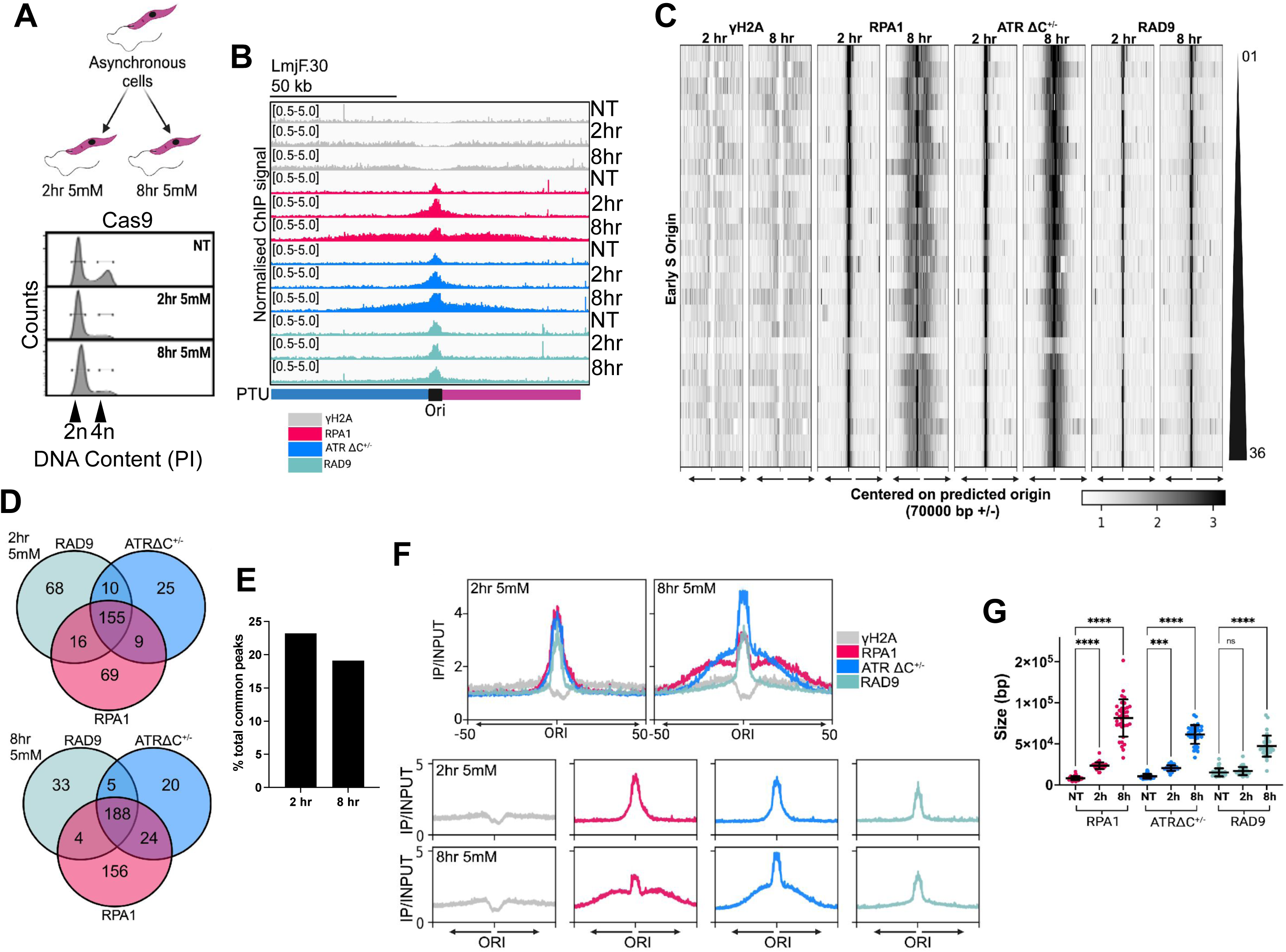
RPA1 accumulates at early origins in synchronised cells. (A) Cartoon illustrating the experiment performed to synchronise cells. (Right) FACs analysis showing the DNA content of *Leishmania* in unperturbed cells and cells collected post exposure to HU (5 mM) for 2 and 8 hrs. (B) RPA1, in ATR competent and ATR deficient cells, and RAD9 co-occupy 155 genomic regions in cells 2 hrs post-HU exposure and 188 genomic regions at 8 hrs post-HU exposure (C). (D) The percentage of the common peaks associated with early origins. (E) Localisation of yH2A, RPA1 and RAD9 in control cells and the location of RPA1 in ATR-deficient cells. ChIP-seq signals shown are mapped to a representative chromosome (LmjF.30). ChIP-seq samples were normalised to their corresponding input controls and shown on a scale of 0.5 - 5.0. Tracks were viewed in IGV. Polycistronic unit negative (blue), polycistronic unit positive (purple), early origin (black). (F) Normalised ChIP-seq signals at 2 and 8 hrs post exposure to 5 mM HU for yH2A, RPA1 (including in ATR-deficient cells) and RAD9 centred on the early origin and plotted +/− 70,000 bp. White = no enrichment. Chromosomes are plotted in order of size. (G) Metaplot analyses depicting enriched signal for yH2A, RPA1 and RAD9 centred on early origin sites +/− 50,000 bp. Below shows the signal for each protein separately. ChIP-seq samples were normalised to their corresponding input controls. (H) The sizes of peaks associated with the early origin were quantified. Significance was calculated using a two-way ANOVA, ns = not significant, *** = < 0.0005, **** = < 0.0001.

By centering the ChIP-seq signals on RPA1-bound regions identified 2 hrs and 8 hrs after HU exposure in RPA1myc cells, we found that the ChIP signals largely remain in proximity to the RPA1-bound regions for 2 hrs, whereas at 8 hrs post HU treatment, we detected a wider enrichment of RPA1 around the origins in WT and ATRΔC+/− cells, while RAD9 remained more proximal to the origins (**Fig. 3B&C**; **Fig. S4C**). We investigated these observations further we examined the locations of the peaks called using MACS2 at 2 and 8 hrs after HU exposure for RAD9 and RPA1. At 2hrs, MACS2 identified 249 peaks in RPA1myc cells, 199 RPA1 peaks in ATRΔC+/− cells, and 249 RAD9 peaks in RAD9myc cells (**Fig. S4D**), after filtering, as described in the methods. The threefold increase in RPA1 peaks in RPA1myc cells compared to untreated controls (**Fig. 2B**) indicates a significant enhancement of RS. In contrast, ATRΔC+/− cells showed a smaller increase (~30%) in RPA1 peaks. After 8 hrs of HU exposure, we observed 373 RPA1 peaks in RPA1myc cells, 237 RPA1 peaks in ATRΔC+/− cells, and 230 RAD9 peaks (**Fig. S4D**). The 50% increase in RPA1 peaks in RPA1myc cells further highlights the active replication stress response in these cells. Despite the rise in RPA1 peaks at both time points, the number of RAD9 peaks only slightly increased (**Figs. S4D and Fig. 2B**).

We then analyzed ChIP signals associated with previously mapped early-S origin regions. After 2 hrs of HU exposure, we identified 155 common RPA1 and RAD9 peaks in both RPA1myc and ATRΔC+/− cells (**Fig. 3D**). After 8 hrs, 188 common RPA1 and RAD9 peaks were observed between these cells (**Fig. 3E**). Approximately 20% of these common peaks colocalized with early-S origins (**Fig. 3D&E**), as nearly all early-S origins were enriched for RPA1 and RAD9, and in ATR-deficient cells. Thus, when compared to untreated cells (**Fig. 2D**), HU treatment induces a broader enrichment of factors to locations beyond the known sites of DNA replication initiation activity which themselves robustly recruit these two replication stress response factors. A more detailed analysis of signal enrichment at the origins (**Fig. 3F-G**) revealed several key observations. First, as HU treatment progressed from 2 to 8 hours, the RPA1 signal significantly extended from all origins in both RPA1myc and ATRΔC+/− cells. This increase in RPA signal on both sides of the predicted origin regions indicates that RS, a known effect of HU exposure, is occurring. Second, RAD9 signal was also present around the early-S origins, but a modest broadening of the RAD9 signalling was only evident after 8 hrs of HU exposure, suggesting this RAD9 signal pattern reflects the parasites response to the accumulation of RPA-bound ssDNA. Finally, whilst untreated cells exhibited no definable yH2A signal accumulation in proximity to the origins (**Fig. 2E**), HU treatment led to a modest increase in signal accumulation distal to both sides of the origins, mirroring that of the RPA1 signal and showing that HU treatment results in DNA damage around the origins (**Fig. F&G**).

In ATRΔC+/− cells, the RPA1 signal was similar to untreated cells after 2 hrs, but at 8 hrs, though we detected a significant increase in peak size, compared with untreated ATR-deficient cells (**Fig. 3G**), the peak appeared more restricted, and the peak sizes were overall shorter than RPA1 origin-associated peaks under the same treatment conditions. In all, we show that though ATR is not crucial for replication initiation, it is likely necessary for maintaining fork progression and/or stability (**Fig. 3F**). Analysis of early origin-associated peaks (**Fig. 3G**) revealed a significant increase in the average size of the RPA1 peaks. After 8 hrs of HU treatment, RPA1 peaks were 10 times broader compared to untreated cells, while RAD9 peaks exhibited a 3-fold increase in width under the same conditions. Altogether, our findings demonstrate that the single origin previously identified by MFA-seq is a major locus of RPA and RAD9 accumulation, suggesting it is the most pronounced site of RS in each chromosome.

### Acute replication stress drives RPA1 accumulation at transcription initiation sites

The chromatin binding profile of RS markers has been utilized previously to study events at unstable genomic regions, including at sites of DNA replication and transcription machinery collisions[13]. Our data above shows that acute HU treatment, which causes cell cycle stalling, leads to a broader localization of RPA1 and RAD9 compared to untreated cells. In unperturbed cells, RPA1 and RAD9 enrichment was observed at early-S origins of replication (48%) and transcription termination sites (49%) (**Fig. 2D, F**); and only 1% of the RPA1 and RAD9 enrichment were found at transcription initiation sites (**Fig. 2D**). Therefore, we next explored the basis of the broader signals detected following acute HU treatment (**Fig.4**). Our analysis of MACS2 peak localisation revealed that 2 hrs of HU exposure resulted in a 30-fold increase in peaks at transcription initiation sites, accounting for approximately 30% of the total common peaks. Additionally, around 26% (41 peaks) were found at both transcription initiation and termination sites, while only about 14% (21 peaks) were at transcription termination SSRs (**Fig. 4A**). A similar pattern emerged at 8 hrs post HU exposure, with ~28% of common peaks at transcription initiation sites, ~23% at both initiation and termination sites, and ~10% at transcription termination-only sites (**Fig. 4B**). Overall, ~56% of the peaks at 2 hrs and ~50% at 8 hrs were associated with transcription initiation. The reduction in common peaks at transcription termination sites after HU treatment likely reflects HU slowing of DNA replication, leading to amelioration of transcription-replication conflicts at these locations. Conversely, the unscheduled interactions between DNA replication and transcription caused by HU-mediated replication inhibition can lead to wider instability. This is illustrated by the increased number of RPA1 and RAD9 peaks across various genomic regions (**Fig. S5**).

**Figure 4.**
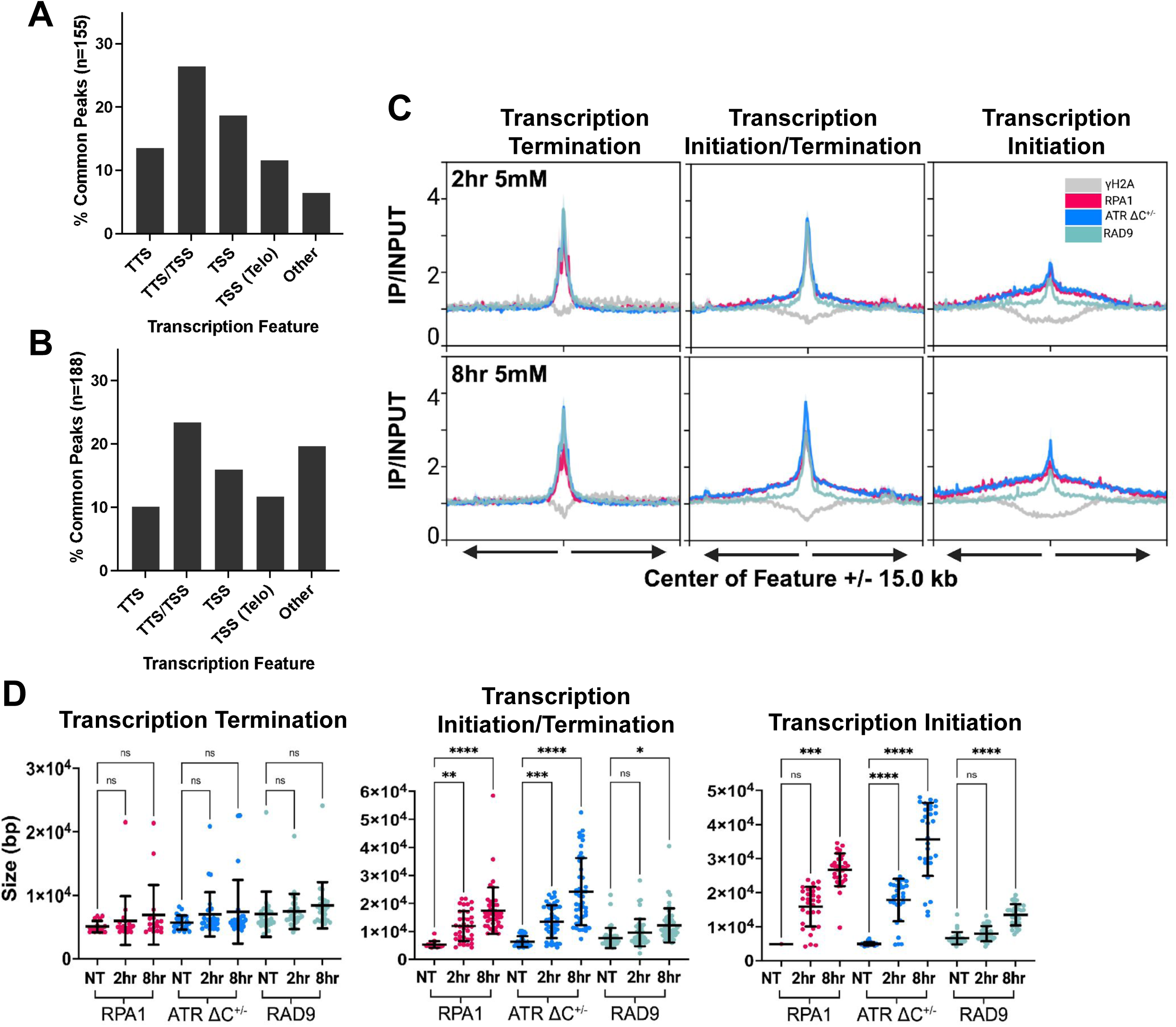
ATR-deficient cells accumulate RPA1 at SSRs associated with transcription initiation. The percentage of the common peaks at 2 hrs (A) and 8 hrs (B) associated with other genomic features. (C) Metaplot analyses depicting enriched signal for yH2A, RPA1 and RAD9 centred on sites enriched for AcH3 (transcription initiation - chromosome cores), BaseJ (transcription termination) and AcH3/BaseJ (transcription initiation and termination) +/− 15 kb at 2 hrs (upper panel) and 8 hrs (lower panel). (D) The sizes of peaks associated with transcription-associated genomic regions were quantified. Significance was calculated using a two-way ANOVA, ns = not significant, *** = < 0.0005, **** = < 0.0001, ** = < 0.005, * < 0.05.

Metaplot analyses (**Fig. 4C**; **Fig. S6**) revealed pronounced peaks of RPA1 and RAD9 at transcription termination sites (BaseJ; left panel), with a modest accumulation of yH2A flanking the center of these regions. However, at transcription initiation sites (AcH3; right panel), the peaks of RAD9 and especially RPA1 extended outward from a modest peak at the center (**Fig. 4C**; **Fig. S6**). Minimal yH2A signal was observed flanking the center of these sites. At sites with both transcription initiation and termination activities (AcH3/BaseJ; middle panel), we observed broader peaks of RPA1 and RAD9. Overall, the RPA1 and Rad9 peak sizes at transcription termination sites (BaseJ) remained consistent across all samples and HU treatment time points (**Fig. 4D**). In contrast, RPA1 peaks at transcription initiation sites (AcH3) showed a significant increase in size at both 2 and 8 hrs of HU treatment compared to unperturbed cells. A similar, though less pronounced, pattern was observed at sites with both transcription initiation and termination activities. RAD9 exhibited a modest increase in peak size at 8 hrs post HU exposure at both AcH3 and AcH3/BaseJ sites. It is noteworthy that in ATRΔC+/− cells, RPA1 peaks were overall broader than those identified in control cells (**Fig. 4D**). This pattern further supports an elevated level of DNA damage and/or increased RS present in these cells. We next examined subtelomeric sites at both 2 and 8 hrs 5 mM HU exposure, revealling an accumulation of both RPA1 and RAD9 at chromosome ends under RS conditions (**Fig. S7A-C**). As these sites are nucleosome depleted [22] we consequently noted a reduction of yH2A in proximity to the subtelomeric region. Little difference was observed between RPA1 signals across ATR competent and ATR-deficient cells.When examining ChIP signal for RPA1, RAD9 and yH2A across subtelomeric-associated transcription initiation sites, we detected an accumulation of RPA1 and RAD9, with a very modest accumulation of yH2A emanating from the region center outwards bidirectionally (**Fig. S7D**). From 2 hrs to 8 hrs the peaks of RPA1 significantly broadened relative to untreated cells and that in ATRΔC+/− cells, RPA1 peaks were overall broader. Though no significant increase in RAD9 peak size was detected, at 8 hrs the peaks we overall broader than 2 hrs and in untreated cells.

Taken together, these findings reveal that while transcription initiation sites are not major stress loci in untreated cells, they exhibit broad and bidirectional accumulation of RPA1 and RAD9 in HU-treated cells, resembling the pattern seen at early-S origins of replication. Under conditions of acute DNA replication stalling, the increased levels of RPA and RAD9 accumulation indicate that sites of transcription initiation are also putative sites of RS, in common with the origins and transcription termination sites. Thus, all PTU boundaries in *Leishmania* appear to be locations of RS.

### During mid S-phase, an origin-like pattern of RPA1 accumulation emerges at transcription initiation sites

Acute RS induced by HU led to a distinct pattern of RPA1 and RAD9 accumulation at transcription initiation sites, similar to that seen at DNA replication origins. We hypothesized that RS might trigger a compensatory adjustment in the density of initiation events. Therefore, we asked what effects are seen when cells are under mild RS conditions that are permissive for DNA replication. *L. major* promastigotes were exposed to a lower dose of HU to induce chronic RS (0.5 mM HU for 8 hrs). We confirmed by FACS analysis an accumulation of cells in S phase (**Fig. 5A**). Under these conditions, where DNA replication from early origins will have progressed, we can explore the dynamics of the cell’s response to RS. We investigated the accumulation patterns of RPA1 in RPA1myc and ATRΔC+/− cells, as well as the RAD9 signal in RAD9myc cells exposed to 0.5 mM HU for 8 hrs (ChIP-seq signal across each chromosome is shown in **Supplementary Data 4**). Under these chronic RS conditions, we observed a poor correlation between the RPA1 signal in RPA1myc cells and ATRΔC+/− cells. In contrast, the correlation between the RAD9 signal in RAD9myc cells and the RPA1 signal in ATRΔC+/− cells remained consistent; together this suggests that the pattern of RPA1 enrichment is altered in ATRΔC+/− cells exposed to chronic HU treatment (**Fig. S8A).**

**Figure 5.**
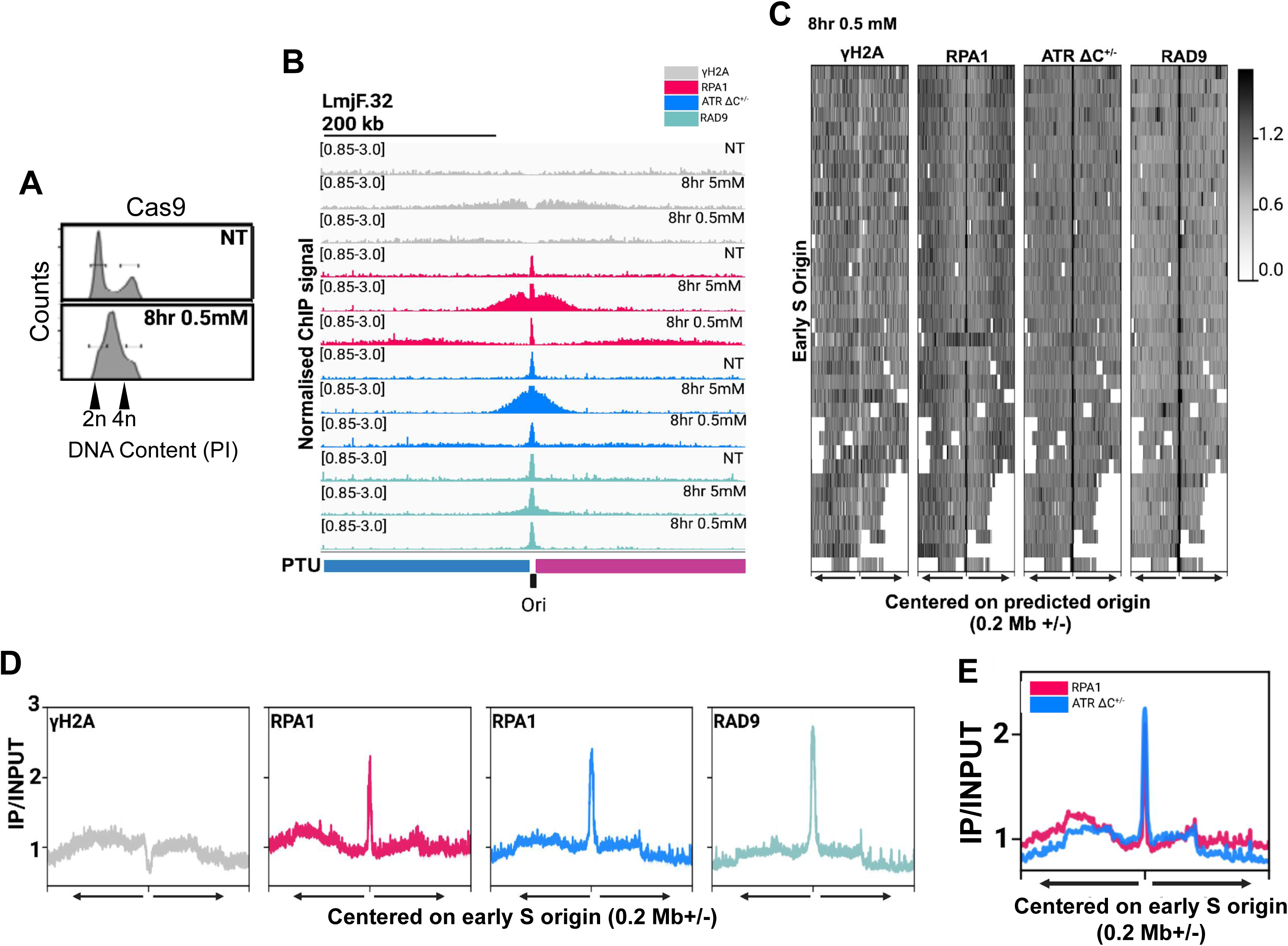
Low dose HU exposure slows DNA replication and reveals ‘hotspots’ of RS at initiation and termination sites for transcription. (A) Representative FACs analysis of the DNA content of promastigotes treated with 0.5 mM HU for 8 hrs. (B) Localisation of yH2A, RPA1 and RAD9 in control cells and the location of RPA1 in ATR-deficient cells. ChIP-seq signals shown are mapped to a representative early origin located on chromosome LmjF. 32. Enriched signals are shown on a scale of 0.85 - 3.0. Tracks were viewed in IGV. Polycistronic unit negative (blue), Polycistronic unit positive (purple), early origin (black). (C) Heatmap of normalised yH2A, RPA1 and RAD9 ChIP signal plotted relative to the early origins (+/− 0.2 Mb). The colour white on the heatmap depicts no binding. (D) Metaplot analyses depicting enriched signals for yH2A, RPA1 and RAD9 at early origins (+/− 0.2 Mb). ChIP-seq samples were normalised to their corresponding input controls. Individual metaplots for each protein. (E) Metaplot analysis depicting enriched signals RPA1 and RPA1 in ATR-deficient cells centred on the early origins +/− 0.2 Mb.

Centering the ChIP-seq signals on early-S origins revealed that, after exposure to 0.5 mM HU, the RPA1 and RAD9 signals were primarily enriched in a relatively narrow region around the origins, as observed in untreated cells. Additionally, the origin-proximal enrichment of RPA1 signal, within 70 Kb upstream or downstream, seen at the 5 mM HU treatment, was not detected in cells treated with 0.5 mM HU (**Fig. S8B**). Instead, these cells exhibited more distal, wider and less intense enrichment of RPA1, within 0.2 Mb from the origin site (**Fig. 5B-D**). Metaplots confirmed these similarities and differences around the mapped origins (**Fig. 5D**) suggesting that the primary effect of chronic HU exposure is accumulation of RPA at RFs that eventually progress bidirectionally from putative replication iniation sites but are likely perturbed due to reduced levels of nucleotides. We found little difference in RPA1 signals between ATR competent and ATR-deficient cells (**Fig. 5E**).

Analysis of ChIP mapping beyond the early-S origins revealed that chronic HU treatment led to widespread accumulation of RPA1 and RAD9 across the genome, with a significant portion of the signal concentrated at PTU boundaries (**Fig. 6**). At transcription termination sites (BaseJ only), sharp peaks of RPA1 and RAD9 were detected, indicating these regions are a consistent hotspot for RS and/or DNA damage (**Fig. 6A and B**). At transcription and termination sites (AcH3/BaseJ), sharp peaks of RPA1 and RAD9 were detected, along with a broadened signal extending towards the base of the peak across all cell lines (**Fig. 6A and B**). Notably, we detected broad peaks of RPA1, RAD9, and yH2A, which were observed upstream and downstream, extending approximately equivalent distances from the transcription initiation sites (AcH3; **Fig. 6A and 6B**). This pattern is reminiscent of the broad peaks identified after acute HU treatment at these AcH3-enriched sites (**Fig. 5**), particularly at early-S origins (**Fig. 4**). Moreover, in ATR-deficient cells, we observed a modest decrease in peak amplitude compared to RPA1 in ATR competent cells, suggesting that ATR could play a role in maintaining RF stability at these sites (**Fig. 6C**). At subtelomeric regions, we identified an accumulation of RPA1 under chronic RS conditions, albeit at a more modest level when compared to acute RS conditions, with a slight reduction in RPA1 signal in ATR-deficient cells (**Fig. S9**). We found little accumulation of RAD9 at these sites and limited yH2A signal, likely due to the reduced nucleosome content within these regions. Thus, though RPA is recruited to ssDNA at these sites, the limited recruitment of RAD9 suggests mild RS exposure may be insufficient to trigger the RSR.

**Figure 6.**
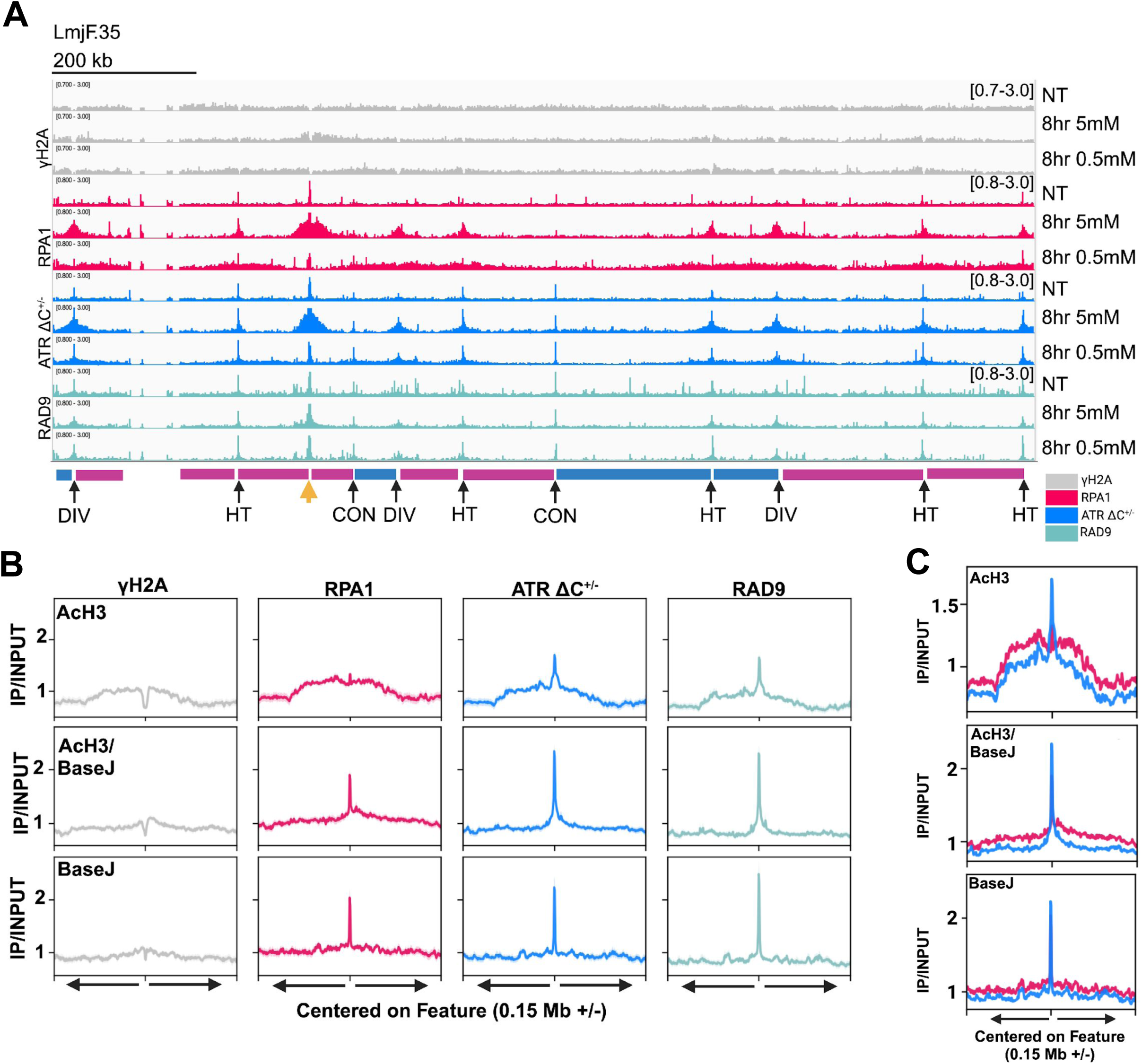
Additional sites of DNA replication initiation co-localise with sites of transcription initiation activity. (A) Localisation of yH2A, RPA1 and RAD9 in control cells and the location of RPA1 in ATR-deficient cells. ChIP-seq signals shown are mapped to a representative chromosome LmjF. 35. Enriched signals are shown on a scale of 0.7 - 3.0 (yH2A) and 0.8 - 3.0 (RPA1 and RAD9). Tracks were viewed in IGV. Polycistronic unit negative (blue), Polycistronic unit positive (purple), early origin (orange arrow). SSR types are labelled below and their locations indicated by black arrows. (B) Metaplot analyses depicting enriched signal for yH2A, RPA1 and RAD9 centred on sites enriched for AcH3 (transcription initiation - chromosome cores), BaseJ (transcription termination) and AcH3/BaseJ (transcription initiation and termination) +/− 0.15 Mb. ChIP-seq samples were normalised to their corresponding input controls. Individual metaplots are shown for each protein. (C) Metaplot analyses depicting enriched signals for RPA1 and RPA1 mapped in ATR-deficient cells centred on different SSR types as detailed in the figure - +/− 0.15 Mb.

Altogether, we suggest that slowing of DNA replication by low levels of HU reveal that SSRs in *Leishmania* are sites for transcription-replication conflicts or replication initiation sites where sensing accumulated ssDNA stretches might be necessary to protect the RF integrity.

## Discussion

The organisation of transcription in *Leishmania*, and indeed in all kinetoplastids, is unique amongst eukaryotes, with virtually all genes transcribed from polycistrons. A consequence of this organisation is that RNA Pol must traverse long distances and, if transcription overlaps with S-phase, it is especially at risk from being impeded by DNA replication. This is the first study that has mapped the consequences of polycistronic transcription in both the absence and presence of RS conditions, uncovering dynamic interactions between DNA replication and transcription in these organisms.

Our global mapping of RPA1 dynamics in the absence of HU highlighted sites of DNA replication initiation during early S-phase as pronounced sites of RS. In the immediate origin centre, we detected sharp peaks of RPA1 and RAD9, indicative of localised DNA damage. Under condition of acute RS, the sharp peaks of RPA1 and RAD9 persisted within the origin centre, and additionally, bi-directional expansions of yH2A and RPA1 emerged, spread outwards from the origin centre. For yeasts and metazoans, and also *Leishmania* (**Fig. 1E**), the origin region is commonly nucleosome depleted [22, 61–63], therefore we were unable to map yH2A signal within the central portion. However, at 8 hrs (5 mM HU) the loss of yH2A immediately proximal to the origin region expanded, indicating the chromatin is more open. These signal patterns began accumulating at 2 hrs post HU exposure, suggesting this response is dynamic. The accumulation of yH2A and RPA1, suggests the presence of expanded ssDNA tracks, and therefore wider stalling of RFs; ssDNA accumulation is commonly a consequence of impaired DNA replication during which the DNA double strand is unwound but uncoupled from DNA synthesis [2].

Under these conditions, RAD9 in *Leishmania* primarily localises to origins, but we noted an expansion of the RAD9 peak, suggesting the recruitment of 9-1-1 to the immediate vicinity of the early-S phase origin likely due to enhanced RS (**Fig. 3**). The recruitment of RSR factors to the origin, even in normally replicating promastigotes, suggests DNA replication initiation here induces especially pronounced levels of RS in *Leishmania*, which is exacerbated after HU treatment. It is likely that the prominence of RSR factors at these single sites in each chromosome reflects their constitutive activation during S-phase and explaining why they are so readily detected by MFA-seq [19, 48]. As these are also sites of both transcription initiation and termination activities, their constitutive use as origins will impose especially focused encounters between the transcription and replication machineries. Though leading DNA strand replication in *Leishmania* moves co-directionally with transcription [19], during lagging strand duplication, head-on collisions between the replisome and the transcription machinery may become more common, inducing DNA breaks; however, we lack strand-specific information from our ChIP-seq analyses to confirm this hypothesis. Efficient, early origins in human cells are associated with enhanced mutagenesis [64] and mutagenic DNA replication initiation coincides with genome evolution, influencing gene expression. As gene expression regulation in *Leishmania* is largely post-transcriptional, the consequences of enhanced RS and likely DNA breaks, within early origins on the regulation *Leishmania* gene expression are unknown. Could origin instability explain in part the frequency of aneuploidy in these parasites, by altering the behaviour of the kinetochore interaction thereby inducing mis-segregation of chromosomes? Enhanced levels of DNA breaks within early origins could increase the level of chromosome fusions, recombination events and genome instability, as reported frequently in DNA repair defective *Leishmania* mutants [65–67]. In the ATR-deficient mutants, RPA1 tracks were highly reduced in length. Though under these conditions, ssDNA becomes exposed, our data suggests that these cells are subject to enhanced fork collapse.

Additionally, under acute HU-induced RS, we uncovered an enhanced accumulation of RPA1 and RAD9 at SSRs, detecting a sharp peak of RAD9 and RPA1 confined to transcription termination sites. The peak amplitude and shape remained largely consistent at 2 and 8 hrs post HU exposure, suggesting these sites are constitute sites of RS, regardless of exposure length. In contrast, at sites of transcription initiation/termination or initiation only, we noted a sharp peak of RPA1 and RAD9 at the centre, which broadened outwards bidirectionally at 8 hrs. At sites of initiation only, the sharp peak at the centre was reduced in amplitude when compared to termination and initiation sites. In both instances, the lack of yH2A signal accumulation and the broad loss of yH2A at the central region outwards suggests a more open chromatin environment (**Fig. 4**). We have not measured DNA replication under these conditions, and no previous work has detected DNA replication activity at all SSRs, even after HU stress[19], but these patterns of RSR enrichment at transcription initiation sites are reminiscent of the accumulation of these factors at the early-S origin, albeit more modestly. Moreover, increasing reduction of yH2A signal at the immediate centre outwards suggests increasingly open chromatin, a feature of sequence consensus-free DNA replication initiation sites[61]. Perhaps, then, these data provide evidence for activation of ‘dormant’ origins at SSRs, which occurs at levels below MFA-seq detection. However, equally these data could be explained by the fact that the activity of RNA polymerase is not inhibited by HU treatment. Thus, under RS the transcription machinery can collide with HU-stalled DNA polymerase wherever it remains in the genome. As transcription start sites are the major sites of RNA polymerase loading, the accumulation of RSR factors to these regions may reflect this. In contrast, at sites of transcription termination, the lack of expanded RPA1 and RAD9 signals suggests localised damage (discussed below in more detail). When we examined the RPA1 signal in ATR-deficient cells, we noted, in contrast to the early-S origin, an increase in peak size at both 2 and 8 hrs relative to RPA1 in ATR competent cells. Thus, ATR may maintain the stability of the sites to prevent excessive ssDNA formation.

Conditions of mild RS (8 hrs, 0.5 mM HU) slow the progression of DNA replication [68]. We did not detect a broad enrichment of yH2A, RPA1 or RAD9 at the early-S origins, only a sharp central peak, suggestive of persistent RS within these sites. Instead, RSR factors accumulate broadly and extend outwards from transcription initiation exclusive SSRs. This pattern was absent at SSRs associated with transcription termination. Instead, sharp peaks of RPA1 and RAD9 were detected, irrespective of HU exposure indicating these sites are constitutive regions of instability and RS. In contrast to initiation sites of transcription, RNA polymerase is unloaded at sites of transcription termination and therefore should not be affected by HU treatment however in human cells, DNA replication terminates at sites of transcription termination[69], perhaps suggest that the instability detected within *Leishmania* could reflects the same process, though more work is required to test this. Given these cells are primarily mid-S phase, the early-S origins have likely fired and progressed, explaining the loss of expanded, bidirectional peaks of yH2A and RPA1 at these sites. RS is associated with the firing of dormant origins in other eukaryotes to ensure replication completion and compensate for stalled RFs if the constitutive origin firing becomes inadequate. Thus, the accumulation of RSR factors largely at transcription initiation sites under chronic RS conditions might provide further evidence that these loci could act as dormant origins, ensuring DNA replication completion under stress. Indeed, evidence for the firing of a single putative dormant origin under RS has been shown in the related pathogen *T. brucei*[70]. One possible benefit of placing a dormant origin at a transcription initiation site is the reduced possibility of head-on collisions with the transcription machinery thereby limiting the formation of DNA damage at these sites[2]. The continuous presence of RAD9 at these sites is indicative of enhanced RS and DNA breaks, suggesting these sites may coincide with enhanced mutagenesis. We found a mild reduction in peak amplitude in ATR deficient cells, suggesting ATR is likely required to maintain the stability of these regions highlight the continual importance of ATR in *Leishmania*.

*Leishmania* subtelomeric duplication appears compartmentalised, operating throughout the cell cycle and driven by RS-associated factors[19]. The clear accumulation of RPA1 and RAD9 signals within subtelomeric regions in both the absence of and during RS is suggestive of a direct relationship between enhanced RS and subtelomeric regions. Moreover, subtelomere-proximal sites of transcription initiation show similar patterns of RPA1 that are suggestive of replication initiation activity. Commonly, expanded gene families like the antigenic genes of pathogens are often located within the subtelomeres[71, 72] thus enhanced RS within subtelomeric regions may diversify multigene families however *Leishmania* subtelomeres are devoid of such genes, thus the importance of enhanced RS within such sites remains unclear.

In conclusion, our study provides the first detailed insights into the dynamics of RSR signalling proteins on the chromatin of *Leishmania*. We identify two major findings. First, the main sites of RS are the boundaries of the PTUs, where RNA polymerase is loaded and unloaded. Second, we observe limited accumulation of RSR factor within the PTUs, even following HU treatment. The alterations in RPA, RAD9, and yH2A signals at SSRs post-HU may suggest increased collisions resulting from DNA polymerase stalling, while RNA polymerase continues its loading and movement; however, these changes might also relate to DNA replication processes, such as the presence of putative ‘dormant’ origins at transcription initiation sites. A possible explanation for having a primary main origin per chromosome could be to minimize collisions between DNA replication and transcription at the few sites where their assembly and disassembly are synchronized. Furthermore, our findings may indicate the existence of previously unrecognized pathways that rapidly resolve conflicts between DNA replication and transcription within PTUs, ensuring continuous RNA polymerase movement and efficient gene expression.

## Materials and Methods

### Lead Contact and Materials Availability

All enquiries regarding additional information/reagents and resources should be directed to and will be fulfilled by the Lead Contact, Prof. Luiz R. O. Tosi. No novel reagents were generated in this work.

### gRNA and donor DNA amplification by PCR

<u>For sgRNA amplification:</u> 0.2 mM dNTPs, 2 µM of sgRNA scaffold and a RPA1-specific forward primer and were mixed in 1× HiFi reaction buffer with MgCl2 and 1 unit Phusion High-Fidelity DNA Polymerase (New England BioLabs), 25 µl total volume. Cycling conditions were as follows: 30 s at 98°C followed by 35 cycles of 10 s at 98°C, 30 s at 60°C, 15 s at 72°C. <u>For donor amplification</u>: 30 ngs of the required pPLOT plasmid [73] was used as a template vector and combined with 0.1 mM dNTPs, 1 µM each of RPA1 donor forward and reverse primers, 1 unit Phusion High-Fidelity DNA Polymerase (New England BioLabs) and 1× HiFi reaction buffer with MgCl_2_. Cycling conditions are as follows: 5 min at 94°C followed by 40 cycles of 30 s at 94°C, 30 s at 60°C, 2 min:30 s at 72°C followed by a final extension step for 7 min at 72°C. Oligonucleotides sequences are provided in **Table S1**.

### Cell Lines and Culture

*L. major* LT252 (MHOM/IR/1983/IR) promastigotes were cultured in M199 medium supplemented with 10 % heat-inactivated fetal bovine serum (FBS; Sigma-Aldrich) at 27 °C as described [31, 74]. Approximately 2×10^7^ exponentially growing Cas9 T7 expressing LT252 cells were transformed with ~4 µg of guide and donor DNA. Cells and DNA were resuspended in 250 μl of phosphate buffer and electroporated using the program X-001 (Amaxa Nucleofector IIb; Lonza). Parasites were recovered in M199 overnight in M199 with 10 % additional FBS. Transformed cells were isolated by limiting dilution in 96 well plates and selected for by the addition of blasticidin. Endogenously tagged RAD9 (RAD9^12myc^) expressing LT252 parasites were generated prior to this study [57]. Parasite proliferation was quantified by assessing cell density every 24 hrs using a Neubauer chamber. Parasites were seeded at a density of 1×10^5^ cells. mL^−1^ and growth monitored for 5 days (120 hrs). Surival curves were performed as follows: cells were seeded at a density of 1×10^5^ cells. mL^−1^ then HU was added at the appropriate concentrations. HU was prepared in M199 fresh at a maximum stock concentration of 200 mM. Surviving cells were then quantified by counting after 96 hrs. Only living cells were considered.

### Genomic DNA extraction and general PCR protocol

For genomic DNA extractions for PCR applications, the Qiagen Blood and Tissue Extraction Kit ® (Qiagen) was used as per the manufacturer’s instructions. PCRs were performed using either Platinum Taq® (Thermo Fisher) or Phusion ® (NEB or Thermo Fisher) as per the manfacturer’s instructions. GC buffer was used for most Phusion® reactions.

### Flow Cytometry

DNA content was assessed by Propidium Iodide (PI) staining and analyzed as described[34]. For experiments involving cell synchronization with HU (Sigma), exponentially growing cells were incubated with 5 mM HU for 8 h. Cells were then washed 1x and recovered in HU-free medium to allow cell cycle progression. To assess the cell cycle under replication stress conditions, exponentially growing cells were treated with 5 mM HU for 2 and 8 h and 0.5 mM HU for 8 and 20 h. Cells were harvested by centrifugation, washed once with 1× PBS and fixed in 30% 1× PBS/70% methanol or ethanol overnight at 4°C. Fixed cells were washed with 1× PBS and stained in 1x PBS containing PI (10 μg. mL^−1^) and RNase A (10 μg. mL^−1^) at 37°C for 30 min in the dark. Flow cytometry data were collected using a BD FACSCanto cytometer (BD Biosciences). Data were analysed using FlowJo. HU was prepared fresh in M199 typically to a maximum concentration of 200 mM.

### Immunofluorescence Analysis

Cells were harvested by centrifugation (1500 *xg*), washed with 1x PBS, fixed with 4% paraformaldehyde for 15 min, then washed with 1x PBS. The cells were settled on Poly-L-Lysine (Sigma) treated slides then permeabilized in 1x PBS, 0.3% Triton X-100 (PBS-T) for 5 min. Cells were blocked with 1% bovine serum albumin (BSA) in PBS-T for 1 hr then incubated with anti-myc antibody at 1:100 dilution in PBS-T, 1% BSA 1 hr. After, cells were washed in 1x PBS then incubated for 1 hr with anti-mouse AlexaFluor488™. DNA was stained with Hoechst 33342 and cells were mounted with ProLong^®^ Gold Antifade Reagent (Thermo Fisher) or DNA was stained using Flouromount-G containing DAPI (Thermo Fisher). Images were captured on a Zeiss Confocal MultiPhoton microscope or using a Leica Fluorescent microscope. Images were prepared and analyzed in Fiji (Image J).

### ssDNA quantification by IFA and flow cytometry

For ssDNA quantification by immunofluorescence analysis, 5×10^5^ cells. mL^−1^ were grown for 16 hrs at 27 °C in 250 µM IdU (protected from light). Cells were then harvested by centrifugation (1400 *xg*, 5 mins) at the time points described in the absence and presence of HU treatment. After, cells were fixed in 4 % paraformaldehyde (500 µL) for 15 mins (agitating, RT), washed in 1x PBS as before then stored in 1x PBS at 4 °C in the dark until required. Cells were settled for 30 mins at RT on poly-L-lysine (Sigma) treated slides, then permeabilized in 1x PBS + Triton X-100 (5 %) for 20 mins at RT. Cells were then washed 5x with 1x PBS then blocked in 1x PBS + BSA (1 %) for 1 hr, RT. After blocking, cells were incubated for 2 hrs RT with anti-BrdU (mouse) antiserum (1:100) diluted in 1x PBS. After, cells were washed 8x in 1x PBS. Cells were then incubated for 30 mins with anti-mouse AlexaFluor488 (1:500) at RT. After, cells were washed 8x in 1x PBS then mounted in a DAPI-containing mounting medium (Southern Biotech). Slides were sealed with nail varnish then stored at 4 °C in the dark until required. To confirm uniform labeling with IdU, controls were denatured using 2N HCL at RT for 30 mins, washed 5x in 1x PBS then neutralized in phosphate buffer 0.2M (0.2M Na_2_HPO_4_, 0.2M KH_2_PO_4_, pH 7.4) for 10 mins. Cells were then washed 5x in 1x PBS then stained as above. Images were captured on a Zeiss MultiPhoton Confocal microscope and the fluorescence intensity quantified using Fiji (Image J).

For ssDNA quantification by flow cytometry, 4×10^5^ cells. mL^−1^ were incubated with 150 µM IdU for 15 hrs at 27 °C (protected from light). After 15 hrs, cells were washed 1x in fresh M199 by centrifugation (1400 *xg* for 10 mins) then resuspended and incubated in M199 containing either 0.5 mM or 5 mM HU. Approximately 2×10^7^ cells were harvested by centrifugation (1400 *xg* for 10 mins) at the stated time points after HU exposure. Everytime point included a corresponding negative control which was not incubated with the primary antibody. Cells were washed 1x in 1x PBS then fixed in 70 % ethanol and stored at −20 °C for at least 16 hrs (protected from light). After fixation, cells were centrifuged at 1400 *xg* for 5 mins then washed in 1x PBS + 1% BSA. The supernatant was discarded and the cell pellet resuspended in 500 µL 1x PBS + 1% BSA containing anti-BrdU (Beckman Dickson; 1:300) for 1 hr RT (cells were placed on a rotor and protected from light). After incubation, cells were washed in 1x PBS + 1% BSA, the supernatant discarded and the pellet resuspended in 500 µL 1x PBS + 1% BSA containing anti-mouse Alexa488™ (1:500). Cells were placed on a rotor and protected from light for 1hr. After, cells were washed in 1x PBS + 1% BSA, then incubated for 30 mins in the dark in 1x PBS containing Propidium Iodide (10 µg. mL^−1^) and RNAse (10 µg. mL^−1^). Cells were then filtered using a 60 µM nylon membrane (Merck Millipore). Flow cytometry data were collected using a BD FACSCanto cytometer (BD Biosciences) and analyzed in FlowJo. The flow rate of the cytometer was kept below 300 cells/second. A minimum of 20000 events were analysed.

### Cell Fractionation

Soluble and chromatin bound proteins were fractionated as described before with the following modifications[31]. Briefly, ~1 × 10^8^ cells were washed with 1x PBS and incubated with buffer A (10 mM Tris-HCl pH 9.0, 100 mM NaCl, 0.1% Triton X-100, 300 mM sucrose, 10 mM MgCl_2_, 10 mM Na_3_VO_4_, 10mM β-glycerophosphate disodium; 3X SIGMAFASTprotease inhibitor cocktail [Sigma]) for 10 min on ice. Samples were then centrifuged (5 min; 3900 ×*g*; 4°C) and the supernatant collected (the soluble fraction; Soluble I). The precipitated insoluble material was treated with buffer A once more, centrifuged as before, then the second soluble fraction (Soluble II) was collected. The remaining pellets were treated with DNase I (Thermo Scientific) (100 units for 5×10^7^ cells) for 1 hr at 37°C. The sample was then centrifuged (5 min; 5000 ×*g*; 4°C), and the supernatant (containing chromatin) collected.

### Immunoblotting

Sodium dodecyl sulfate-polyacrylamide gel electrophoresis (SDS-PAGE) was used to resolve proteins that were then transferred to Polyvinylidene difluoride (PVDF) membrane or nitrocelulose membrane. Before probing for specific proteins, membranes were blocked with 10% (w/v) non-fat dry milk in phosphate-buffered saline (PBS). ECL Prime Western Blotting Detection Reagent (GE Life Sciences) was used for band detection as visualized with Hyperfilm ECL (GE Life Sciences) or using a GE ImageQuant LAS-4000 Multi-Mode Imager.

### Southern Blotting

For Southern blotting, DNA was extracted as described in [74]. Genomic DNA was initially digest with NotI and the digested products separated by gel electrophoresis using the following conditions (0.6 % agarose in 1x TAE buffer, 20 Volts for 16 hrs). DNA was transferred to a Hybond-N+ membrane (GE Life Science). Probes were generating via a PCR: probe for ATR gene (419pb, FW-GACGAGCTGGACCGTTACAT, RV-CTCGCCTCGTAGATGCTCTG) and one for the mutated allele, in this case regions from the Puromycin gene (426bp, FW-GTCACCGAGCTGCAAGAACT, RV-GTCCTTCGGGCACTCGAC). The hybridisation was performed at 65 ^0^C. For probe labelling, the AlkPhos Direct Labelling and Detection system was as per the manufacturer’s instructions and signal visualised using CDP-Star developing reagent (GE Life Sciences).

### Chromatin Immunoprecipitation (myc tagged proteins)

ChIP for RPA1 and RAD9 was performed as detailed[26] with the following modifications. Exponentially growing promastigotes were harvested in the presence or absence of HU treatment. Cells were fixed in formaldehyde (1 %; Sigma) for 40 mins (agitating), quenched in 125 mM Glycine (Sigma) for 5 mins (agitating) at room temperature then samples centrifuged at 1400 *xg* for 5 mins. Next, the supernatant was discarded, the pellets resuspended in 1x PBS/125 mM Glycine solution then left at room temperature (with agitation) for 5 mins. Samples were centrifuged and washed as before then the pellets lysed in Lysis Buffer 1 (HEPES-NaOH 50 mM pH 7.5, NaCl 140 mM, EDTA 1 mM, glycerol 10 %, NP-40 0.5%, Triton X-100 0.25%, 1x protease inhibitors [SIGMAFAST™ Protease Inhibitor Cocktail Tablets, Sigma]) for 10 mins (agitating) at 4 °C. The samples were centrifuged again (at 4 °C) then resuspended in Lysis Buffer 2 (Tris-HCl 10 mM pH 8.0, NaCl 200 mM, EDTA 1 mM, EGTA 0.5 mM, 1x protease inhibitors) for 10 mins (agitating) at room temperature. Next, samples were centrifuged as before then Lysis Buffer 3 (Tris-HCl 10 mM pH 8.0, NaCl 100 mM, EDTA 1 mM, EGTA 0.5 mM, Na-Deoxycholate 0.1 %, N-lauroylsarcosine 0.3 %, 1x protease inhibitors) added and the samples kept on ice prior to sonication. Sonication was performed using a QSonica Sonicator Model Q125 under the following conditions (30 cycles, 45 seg ON/ 59 seg OFF pulsing, amplitude 60 %). Triton X-100 (1.1 %) was then added and the samples centrifuged at 10 000 *xg* for 2 mins (4 °C). For immunoprecipitation of myc bound chromatin, Dynabeads™ (Sheep anti-mouse IgG, ThermoFisher) were washed with blocking buffer (1x PBS, BSA 0.5 %, EDTA 2 mM) then 5 µL of anti-myc antiserum (in blocking buffer) and the sonicated samples added to the beads and left rotating for 5 hrs (4 °C). After, the beads were washed in TSE 150/500 buffer (Triton X-100 1%, SDS 0.1 %, EDTA 2 mM, Tris-HCl 20 mM pH 8.0, NaCl 150 mM / 500 mM, 1x protease inhibitors), then LiCl wash buffer (LiCl 0.25 M, NP-40 1 %, Na-Deoxycholate 1 % EDTA 1 mM, Tris-HCl 10 mM pH 8.0, 1x protease inhibitors) and finally in TE buffer (Tris-HCl pH 8.0, EDTA, 1X protease inhibitors). After TE buffer was removed, the beads were resuspended in 200 µL Elution Buffer (Tris-HCl 50 mM pH 8.0, EDTA 10 mM, SDS 1 %) and incubated at 65 °C for 30 mins (with agitation every 5 mins). Next, the elute was separated from the beads and incubated at 65 °C overnight to reverse crosslinks. After 4 µL of RNAse A (20 mg/mL) was added for 2 hrs at 37 °C then 4 µL of Proteinase K (20 mg/mL) was added for 2 hrs at 55 °C. DNA was purified using the Illustra™ GFX™ PCR DNA and Gel Band Purification Kit (GE Healthcare) as per the manufacturer’s instructions. DNA quantification was performed using a Qubit^®^ 2.0 Fluorometer (Invitrogen).

### Chromatin Immunoprecipitation (yH2A)

Suboptimal immunoprecipitation of yH2A occurred using the above protocol, therefore, a modified protocol based Wedel and Siegel, 2017 [75] was used. Briefly, cultures were prepared and treated with hydroxyurea as described above. Cells were harvested by centrifugation at 1000 *xg* for 10 mins, the resulting pellet re-suspended in M199 medium (lacking serum) then crosslinked in using formaldehyde (50 mM HEPES-NaOH pH 7,5, 100 mM NaCl, 1 mM EDTA, 0,5 mM EGTA, 11% (v/v) formaldehyde) for 20 mins (with agitation). The reaction was quenched by the addition of 125 mM Glycine (Sigma) for 5 mins at room temperature (while agitated). After, samples were centrifuged at 2000 *xg* for 20 mins (4 °C), the supernatant removed, and the pellets washed by centrifugation in 1x PBS. Cells were then resuspended in permeabilization buffer containing protease inhibitors (100 Mm KCl, 10 mM Tris pH 8.0, 25 mM EDTA pH 8.0, 1X protease inhibitors [SIGMAFAST™ Protease Inhibitor Cocktail Tablets, Sigma]). Fifty microlitres of Digitonin (4 mM) were added and the samples incubated for 15 mins (with agitation) at room temperature. Samples were subject to centrifugation at 1400 *xg* for 5 mins (4 °C), the supernatant discarded, the pellet resuspended in ice cold NP-S buffer (0.5 mM Spermidine, 0.075% (v/v) IGEPAL, 50 mM NaCl, 10 mM Tris-HCl pH 7.5, 5 mM MgCl_2_, 1 mM CaCl_2_, 1x protease inhibitors) and the samples vortexed. Next, samples were centrifuged at 1400 *xg* for 5 mins (4 °C), the previous step repeated then the pellets resuspended in ice cold NP-S containing 4U MNase (New England Biolabs) for 10 mins at 25 °C. After 0.5 mM EDTA was added, the samples centrifuged at 10 000 xg for 10 mins (4 °C) and the supernatant with the mononucleosome fraction transferred to a new tube. The remaining pellets were resuspended in ice cold NP-S buffer as before containing 2 ul 10 % SDS then subject to sonication, on ice, for 10 cycles of 30 second on/30 second off pulses (20 % amplitude; QSonica Sonicators Model Q125). Next, samples were centrifuged at 10 000 *xg* for 10 mins (4 °C), the supernatant removed and combined with the mononucleosome fraction. The immunoprecipitation was performed as detailed for RPA1 and RAD9. Dynabeads™ M-280 Sheep anti-Rabbit IgG (ThermoFisher) were used to capture 5 µL of anti-yH2A antiserum raised against the *T. brucei* peptide [51] for each sample.

### Library Preparation

ChIP samples were prepared for next-generation sequencing using the TrueSeq ChIP Library Preparation Kit (Illumina) at the University of Glasgow. Agencourt AMPure XP beads (Beckman Coulter) were used to size select for 300 bp fragments (including adaptor sequence). All samples were sequenced at Glasgow Polyomics (University of Glasgow) in a NextSeq 500 Illumina machine. Paired end reads were generated. For the second replicate of the yH2A ChIP-seq, single-ended libraries were prepared by BGI and sequenced using their DNB-seq technology, with ~20 million single-end reads per sample (Phred +33). ChIP-seqs of RPA1, yH2A and RPA1 in ATR-deficient cells were performed as biological duplicates. ChIP-seq of RAD9 was performed once and a ChIP-seq of a second member of the 9-1-1 complex (Hus1), under the same conditions, is provided as supplementary data to support global distribution pattern of RAD9 (**Supplementary Data 2**).

### ChIP-seq Data Analysis

Sequence read quality was examined first by FastQC[76]. After, adapter sequences were trimmed off using TrimGalore (default setting) and the trimmed reads mapped to the *Leishmania* LmjF.b41 reference genome (TriTrypDB.orf) using Bowtie2[77] -- verysensitive --). Input and IP samples were normalised using the SES method by the DeepTools[78] bamCompare tool and the fold-change calculated as a ratio of the reads. Duplicate sequences were ignored, and ‘Shade-regions’ were removed from the downstream analysis (regions of genes with high copy number as described [22]). ChIP tracks were visualised in IGV v2.14[79]. Peaks were called using MACS2[80] broad peak calling settings with the following conditions: -- gsize 32900000, -- extsize 200, -- nomodel, --shift 250, -- qvalue 0.01. Metaplots and heatmaps were generated using DeepTools ComputeMatrix and the plotProfile or plotHeatmap function using the normalised ratio files. K-means clustering was performed as described in the corresponding figure legends. For Z-Scores, values were calculated per chromosome in RStudio using a custom script (available upon request). Coverage across all chromosomes is shown as supplementary data (**Supplementary Data 3-5)**.

### Data Presentation, Statistical Analysis and Software Usage

Figures were organized in MS PowerPoint ®. Data was visualized using Prism v10 (GraphPad), R and RStudio (www.r-project.org; R Development 2010). ChIPseq data was examined in the IGV. All statistical analysis was performed in R or using Prism. Statistical tests used are as described in the corresponding figure legends. MS Excel (Microsoft®) was used to prepare data tables. Some sequence data analysis was performed using the Galaxy server (usegalaxy.org/usegalaxy.eu [81]), with the remainder performed on internal servers. Flow cytometry data was analyzed using FlowJo v.10 (https://www.flowjo.com). Figures were generated using BioRender. All raw data underlying each figure (where applicable) is shown in Table S2. Custom scripts for data analyses will be provided upon reasonable request.

### Datasets used in this study

Raw sequence reads for KKT1 ChIP were acquired from [20]. BaseJ and AcH3 coordinates were provided by JD Damasceno and originally acquired from [28, 29, 82]. Nucleosome data was acquired from Lombrana et al. 2016[22]. All ChIPseq data generated in this study is accessible via the European Nucleotide Archive (ENA), accession number X. This study would not have been possible without TriTrypDB.org [83, 84].

## Author contributions

SV, JAB, MBS, GLAS, BG generated and analysed data. KC and JDD provided bioinformatics support and genome files. RMC and LROT secured funding and directed the study. JAB and LROT wrote the manuscript. All authors contributed to the editing of this manuscript.

## Acknowledgements

We thank members of the Tosi and McCulloch labs for their insightful discussions and critical reading of the manuscript.

## SUPPLEMENTARY DATA

**Supplementary Data 1:** HUS1 ChIP signal in untreated cells and following HU treatments

**Supplementary Data 2:** Whole chromosome plots of ChIP signal in untreated cells

**Supplementary Data 3:** Whole chromosome plots of ChIP signal under acute RS

**Supplementary Data 4:** Whole chromosome plots of ChIP signal under chronic RS

**Table S1:** Oligonucleotide sequences used in this study

**Table S2:** Original data underlying graphs in Figure 1 and the corresponding supplementary figures S Figure 1 and S Figure 2.

## Supplementary Figures

**Figure S1.**
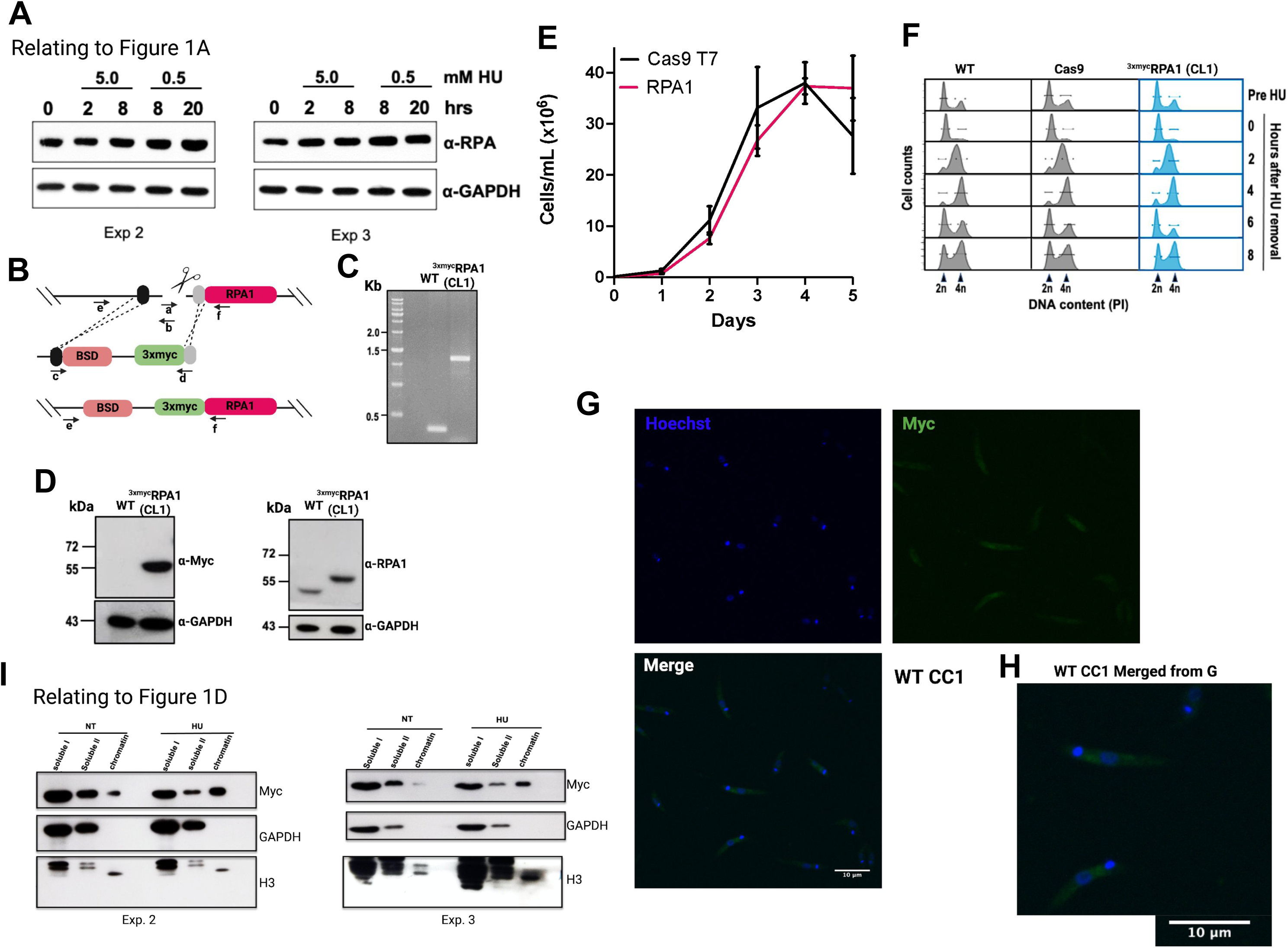
Endogenous tagging LmRPA1 does not affect cell cycle progression. (A) Repeats of immunoblots relating to figure 1A showing RPA1 and yH2A signal after exposure to HU at two concentrations. Graph below shows the RPA1 signal quantified relative to the loading control (displayed in arbitrary units; A.U). (B) Schematic representation of the method used to endogenously tag RPA1 with 3 myc epitopes at the N-terminus of the protein using CRISPR/Cas9 genome engineering. (C) PCR confirmation of tag insertion into the endogenous locus. Immunoblot analyses confirming tag expression are shown below using anti-RPA1 (D) and anti-myc. (E) Growth of the endogenously tagged RPAmyc cells were compared to Cas9 T7 expressing control cells. Growth was assessed every 24 hrs by quantifying cell density. Error bars = +/− SD, n = 3 independent experiments. (F) Cell cycle progression was blocked using 5.0 mM HU for 8 hrs and RPAmyc, WT CC1 and Cas9 T7 cells reseeded in HU-free medium. Cells were collected at regular intervals after HU release as shown; DNA content was examined by flow cytometry. Each histogram represents data from 50,000 events. PI = propidium iodide, n = 3 independent replications/cell line. FACs data was analysed using FlowJo. (G) Representative field of view images from untagged control cells. DNA and kDNA were stained using Hoechst (blue), and anti-myc antiserum used to detect my signal (green). Images were taken on a Zeiss Multiphoton confocal microscope, scale bar = 10 µm. Enlarged and merged images are shown in H. Scale bar = = 10 µm. (I) Repeats of immunoblots from the chromain fractionation shown in relating to figure 1D.

**Figure S2.**
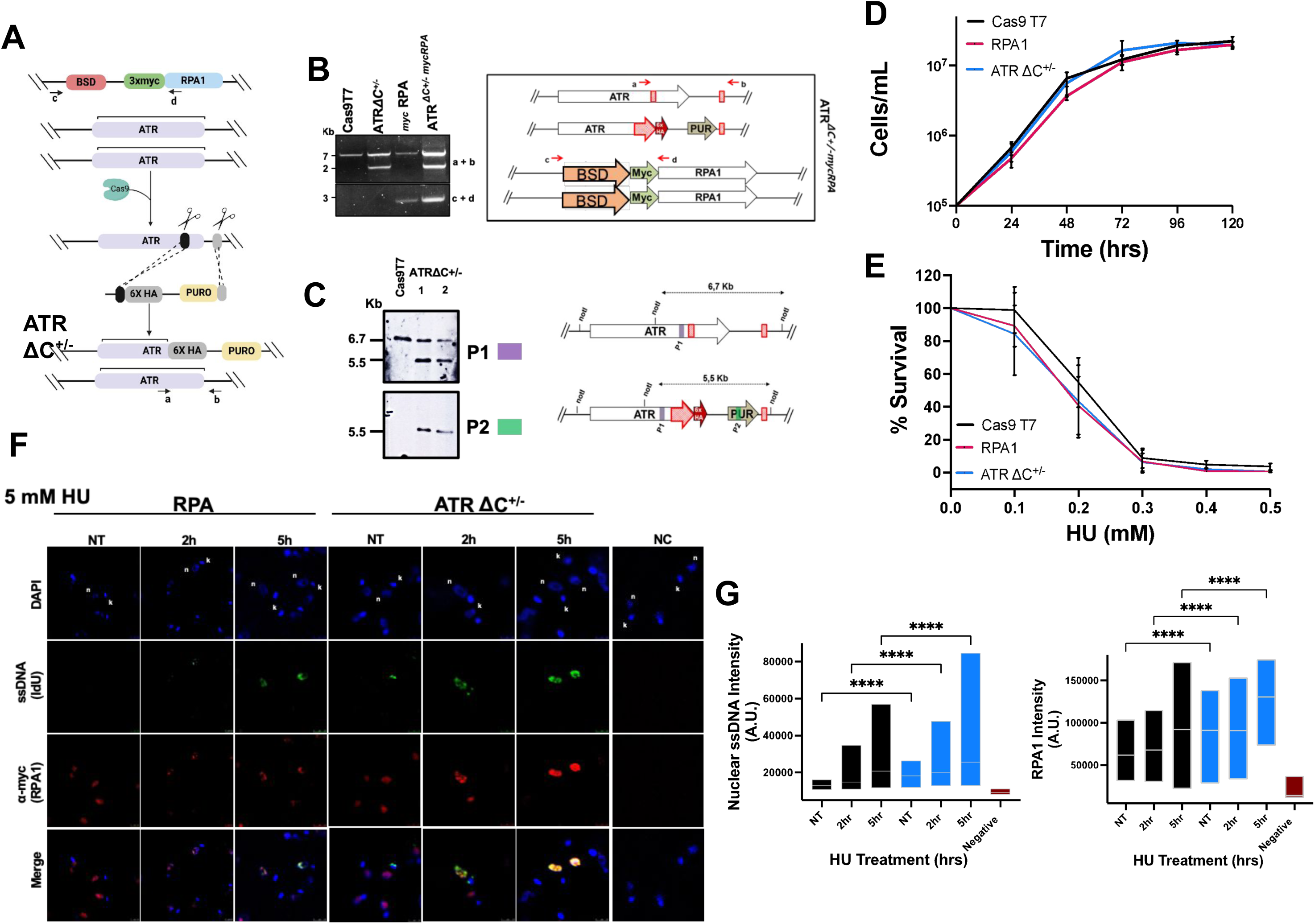
ATR-deficiency enhances RS in *Leishmania* promastigotes. (A) Schematic representation of the method used to truncate one allele of ATR in parasites expressing RPA1 fused to 3 *myc* epitopes at the N-terminus of the protein using CRISPR/Cas9 genome engineering. (B) PCR confirmation of truncation of one allele of ATR and schematic of primer locations used to confirm truncation (right). (C) Southern blotting reveals the cells possess one WT allele (6.7 kb) and one truncated allele (5.5 kb). Right shows schematic representing the Southern blotting strategy to confirm truncation of one allele of ATR. Location of probe 1 = grey square, location of probe 2 = green square. Cas9 T7 and ATRΔC^+/−^ cells were digested with NotI for blotting. (D) Growth of the endogenously tagged RPAmyc cells were compared to Cas9 T7 and ATRΔC^+/−^ cells. Growth was quantified every 24 hrs by examining cell density. Error bars = +/− SD, n = 3 independent experiments. (E) Survival curve examining the percentagem (%) survival of Cas9 T7, RPAmyc and ATRΔC^+/−^ cells after exposure to diferente concentrations of HU. Cells were counted 96 hrs after HU addition. Error bars = +/− SD, n = 3 independent experiments. (F) Representative images of ssDNA signal (green) and anti-myc signal (red; RPA1) in promastigotes in the absence and presence of HU treatment (5 mM; 2 hrs and 8 hrs) in control cells and cells expressing the truncation of one allele of ATR. Nuclear and kinetoplastid DNA is visualised by DAPI incorporation (blue). Images were captured on a Zeiss Multiphoton confocal microscope. Images were processed in Fiji (ImageJ) and colours were enhanced for improved visualisation. Representative of n = 2 independent repeats. Negative control cells from the experimente are show in the last panel (NC, negative control). (G) Quantification of nuclear ssDNA intensity (left) and RPAmyc (right) intensity in RPAmyc and ATRΔC^+/−^ cells. Intensity is expressed as arbitrary units (AU). Statistical significance was assessed using a Kruskal-Wallis test followed by Dunns Multiple comparisons. Representative of n = 2 independent repeats. Signal intensity was quantified using Fiji (Image J) and untagged Cas9T7 cells were used as negative controls.

**Figure S3.**
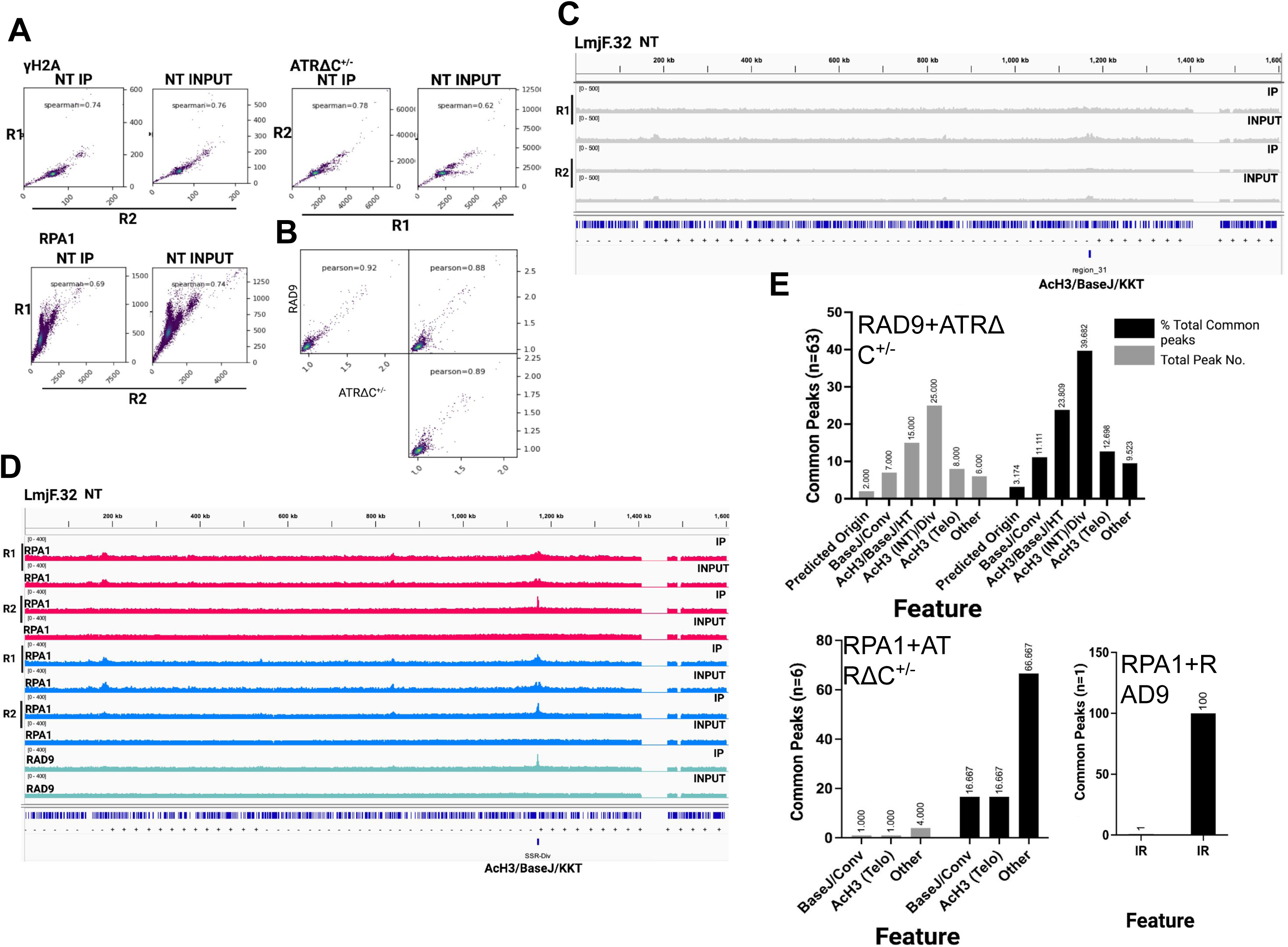
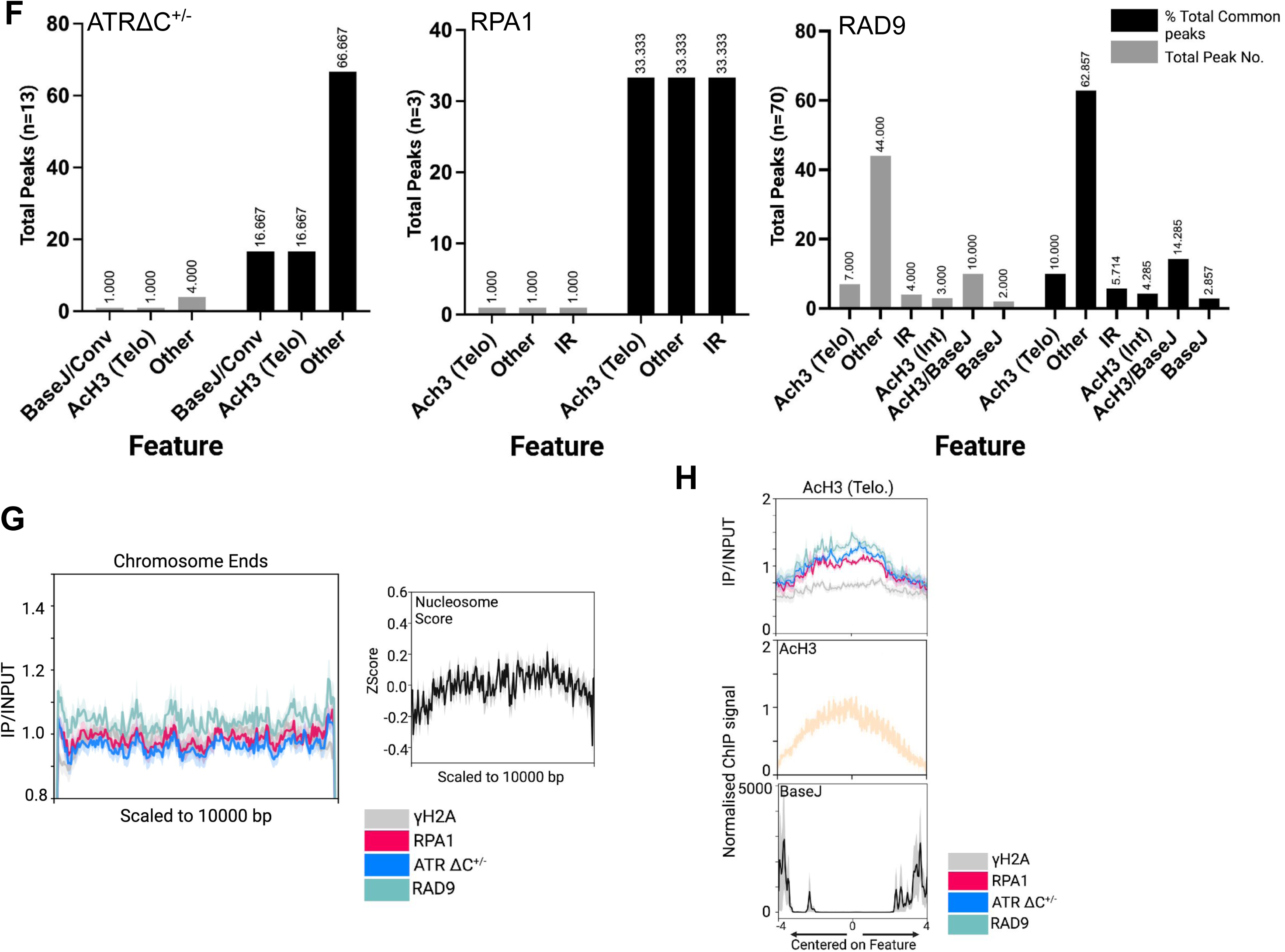
Global distribution of yH2A, RPA1 and RAD9 in unperturbed cells and in unperturbed cells with one allele of ATR truncated. (A) Spearman coefficient correlation analysis to assess the global comparison of ChIP-seq signal between biological duplicates for yH2A and RPA1 (in ATR-competent and ATR-deficient cells) revealed a strong correlation between replicates (B) Pearson coefficient correlation analysis to assess the global comparison of RPA1 ChIP-seq signal between RPA1 signal in ^3myc^RPA1 and ^3myc^RPA(ATRΔC^+/−^) cells and RAD9^12myc^ ChIP-seq signal in unperturbed cells uncovered strong correlations between all three ChIP-seq datasets (C) Normalised coverage files were prepared using bamCoverage (Deeptools) for yH2A signal from immunoprecipitated and input samples taken from unperturbed cells. Replicates (R1 & R2) for both RPA1 ChIP-seq datasets are shown separately. ChIP-seq signals shown are mapped to a representative chromosome (LmjF.32). Samples are shown on a scale of 0-400. Tracks were viewed in IGV. CDSs are shown as blue lines. The early origin is labelled (D) Normalised coverage files were prepared using bamCoverage (Deeptools) for RPA1 and RAD9 signal from immunoprecipitated and input samples taken from unperturbed cells. Replicates (R1 & R2) for both RPA1 ChIP-seq datasets are shown separately. ChIP-seq signals shown are mapped to a representative chromosome (LmjF.32). Samples are shown on a scale of 0-400. Tracks were viewed in IGV. CDSs are shown as blue lines. The early origin is labelled (E) The genomic features associated with common peaks are shown as a percentage of the total number of common peaks (black) and as the number of peaks (grey). (F) The genomic features associated with unique peaks of RPA1, RPA1 (ATR-deficient) and RAD9 shown as a percentage of the total number of common peaks (black) and as the number of peaks (grey). (G) Metaplot analysis depicts enriched signals for yH2A, RPA1 and RAD9 in unperturbed cells within subtelomeric sites. Signals were scaled to 10,000 bp. ChIP-seq samples were normalised to their corresponding input controls. (Left) Metaplot analysis depicting nucleosome enrichment at the subtelomeric regions of chromosomes. Signal is shown as a normalised Zscore. Signal is scaled to 10,000 bps. (H) Metaplot analysis depicting enriched signal for yH2A, RPA1 and RAD9 at 2 hrs and 8 hrs centred on sites enriched for AcH3 within subtelomeric regions enriched for AcH3 (AcH3 Telo) +/− 4 kb.

**Figure S4.**
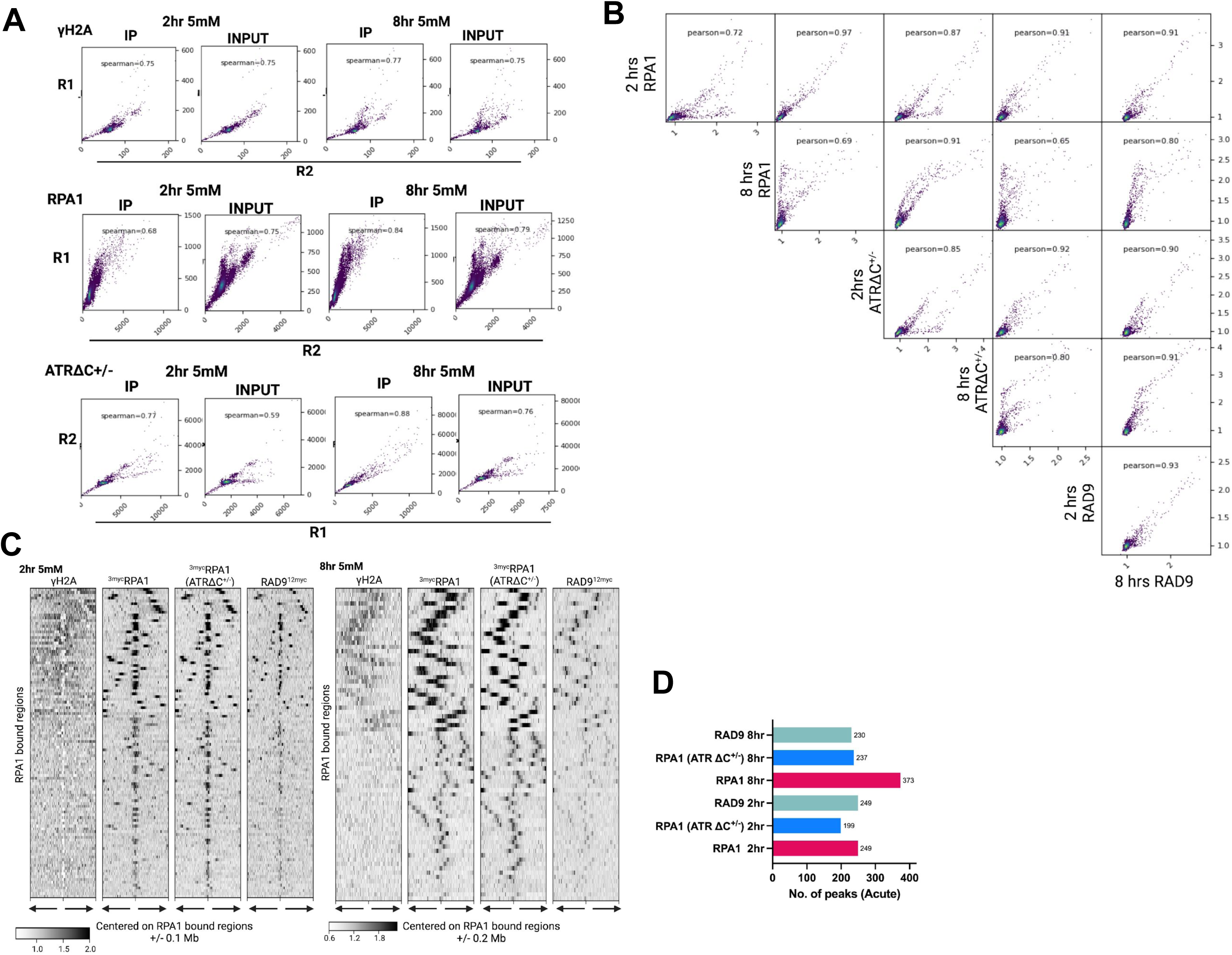
Under acute RS, RSR factors accumulate around sites of DNA replication initiation. (A) Spearman coefficient correlation analysis to assess the global comparison of ChIP-seq signal between biological duplicates for yH2A and RPA1 (in ATR-competent and ATR-deficient cells) revealed a strong correlation between replicates at 2 hrs and 8 hrs post exposure to 5 mM HU. (B) Pearson coefficient correlation analysis to assess the global comparison of RPA1 ChIP-seq signal between RPA1 signal in ^3myc^RPA1 and ^3myc^RPA(ATRΔC^+/−^) cells and RAD9^12myc^ ChIP-seq signal at 2 hrs and 8 hrs uncovered strong correlations between RPA1 signal in ATR-competent and ATR-deficient cells and between RPA1 in both cell lines and RAD9 (C) RPA1-bound sites identified using MACS2 were used to plot the signals for yH2A, RPA1, RPA1 (in ATR-deficient cells) and RAD9 at 2 (left heatmap) and 8 hrs (right heatmap) post HU exposure. The enriched signals were centred on these sites and signal plotted +/− 0.1 Mb (2 hrs) and +/− 0.2 Mb (8 hrs) up- and down-stream of the regions. The colour white on the heatmap shows no binding (D) The total number of peaks for RPA1 and RAD9 identified using MACS2 at 2 and 8 hrs post HU exposure. Peaks were filtered for quality (> 20) and removed if associated with regions of high coverage (‘shade regions’).

**Figure S5.**
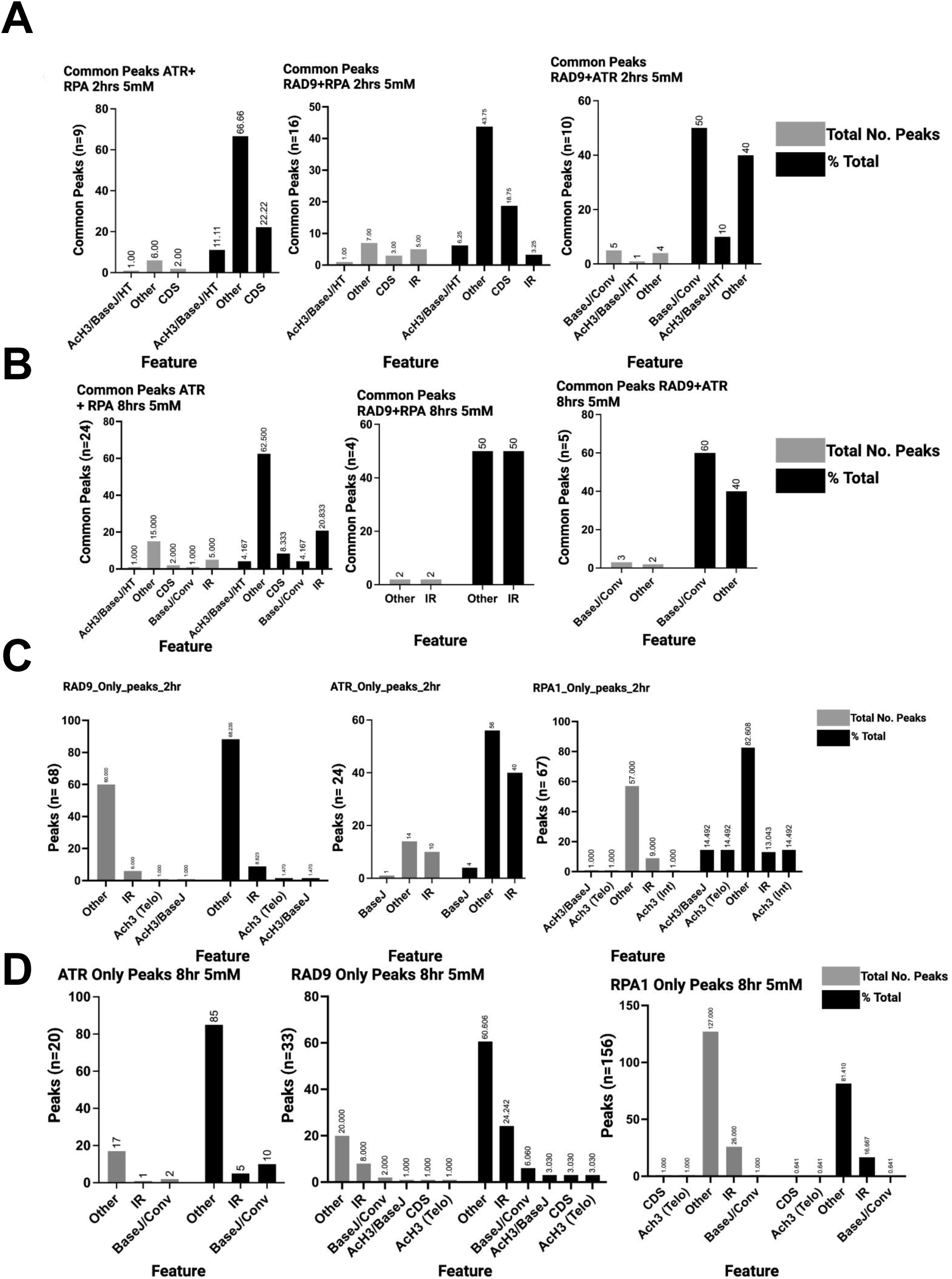
Genomic features associated with peaks of RAD9 and RPA1 under acute RS treatment. The genomic features associated with common peaks at 2 hrs (A) and 8 hrs (B) post HU treatment are shown as a percentage of the total number of common peaks (black) and as the number of peaks (grey). The genomic features associated with unique peaks of RPA1, RPA1 (ATR-deficient) and RAD9 at 2 hrs (C) and 8 hrs (D) are shown as a percentage of the total number of common peaks (black) and as the number of peaks (grey).

**Figure S6.**
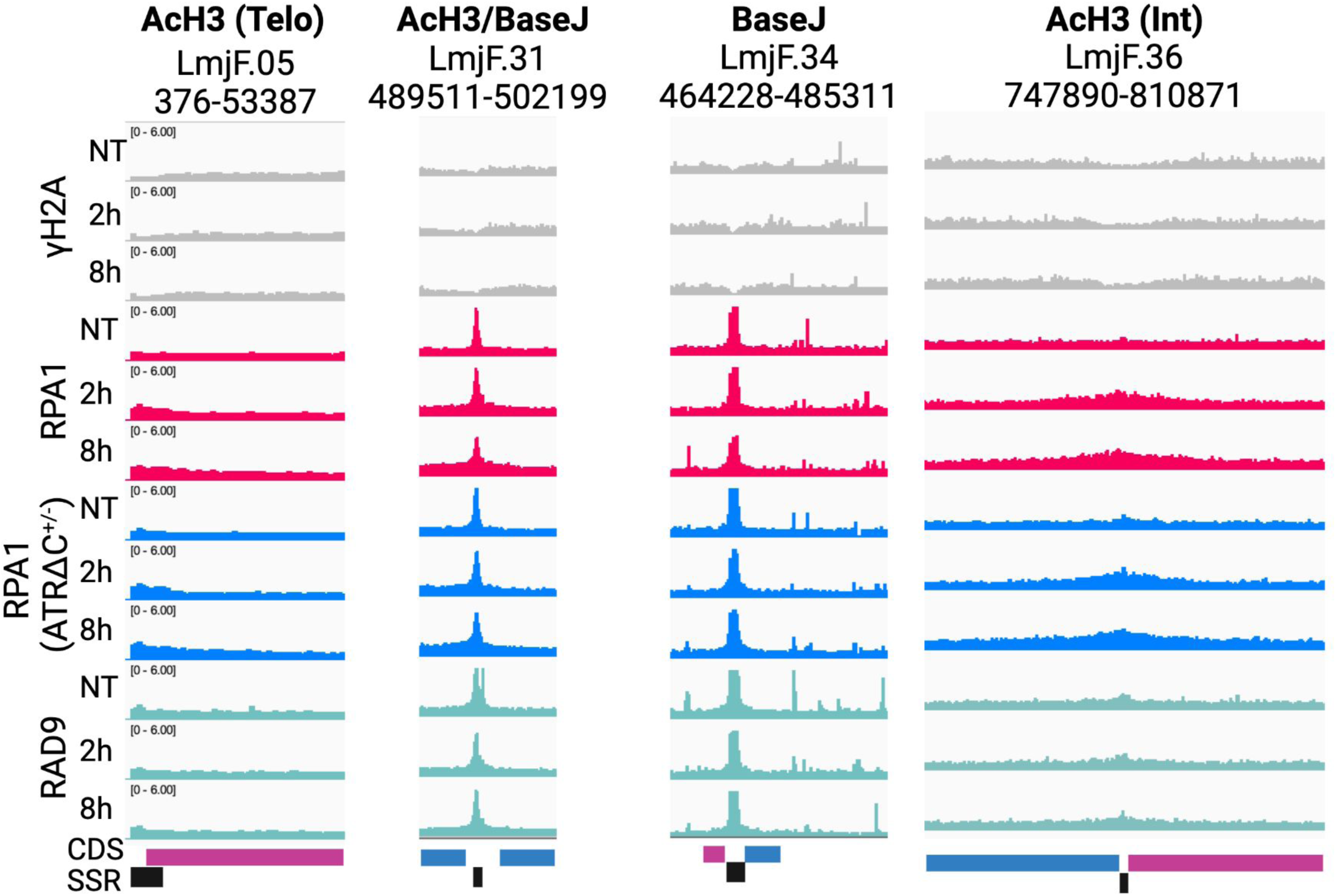
Pattern of ChIP-seq signal enrichment at different SSR types in promastigote cells treated with 5 mM HU. Localisation of yH2A, RPA1 and RAD9 in control cells and the location of RPA1 in ATR-deficient cells in unperturbed cells and cells at 2 hrs and 8 hrs post HU exposure. ChIP-seq signals shown are mapped to representative SSRs on various chromosomes as detailed in the figure. ChIP-seq samples were normalised to input. Scale = 0.0 - 6.0. Tracks were viewed in IGV. Polycistronic unit negative (blue), Polycistronic unit positive (purple), SSR (black box).

**Figure S7.**
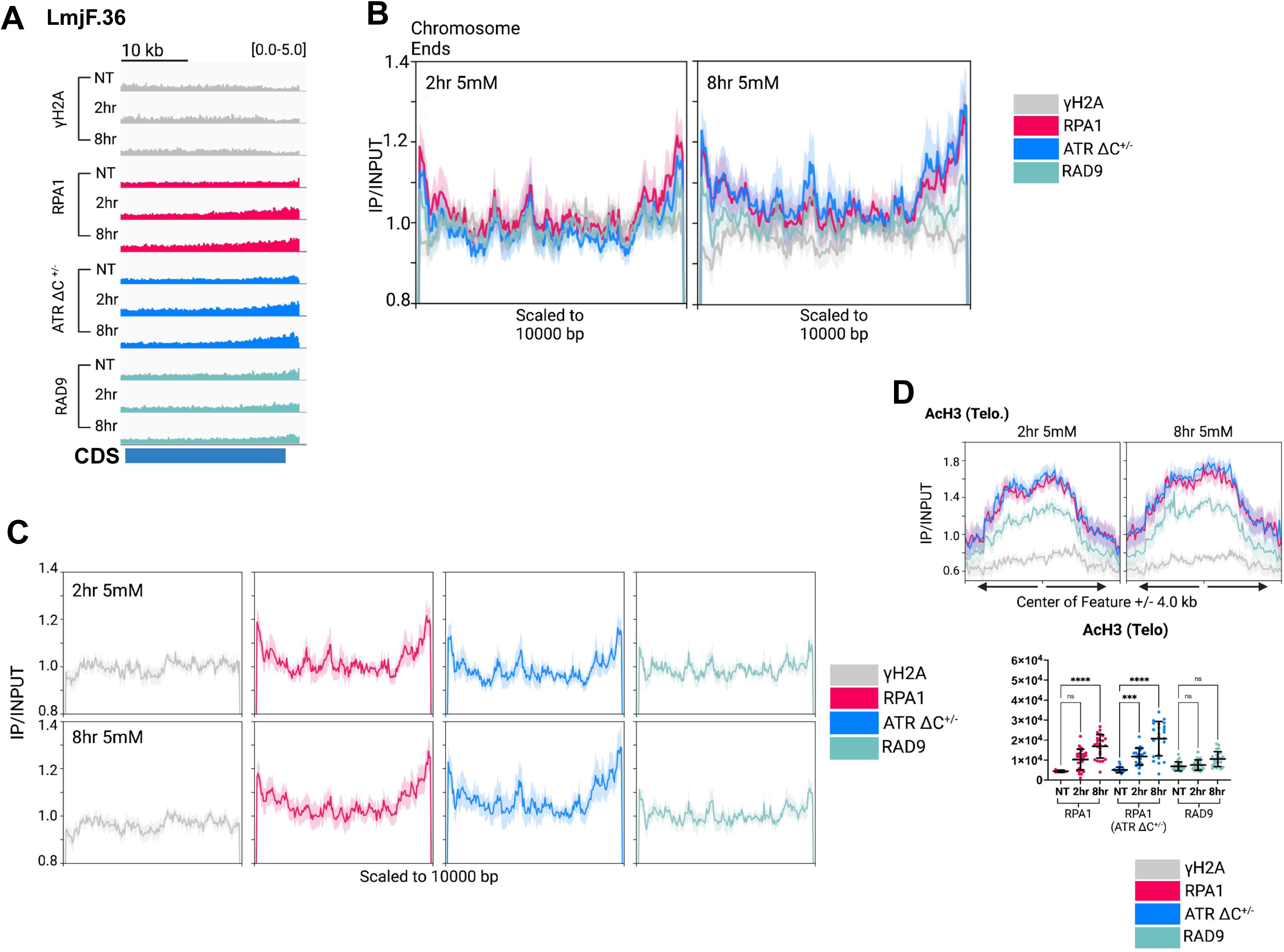
RS associated factors accumulate at subtelomeric sites in cells treated with HU. (A) Localisation of yH2A, RPA1 and RAD9 in control cells and the location of RPA1 in ATR-deficient cells. ChIP-seq signals shown are mapped to a representative subtelomeric region on chromosome LmjF. 36. ChIP-seq samples were normalised to their corresponding input controls and shown on a scale of 0.0 - 5.0. Tracks were viewed in IGV. Polycistronic unit negative (blue). (B) Metaplot analyses depicting enriched signals for yH2A, RPA1 and RAD9 within subtelomeric sites. Signals were scaled to 10,000 bp. ChIP-seq samples were normalised to their corresponding input controls. Individual metaplots for each protein are shown in (C). Signal is scaled as for B. (D) Metaplot analysis depicting enriched signal for yH2A, RPA1 and RAD9 at 2 hrs and 8 hrs centred on sites enriched for AcH3 within subtelomeric regions enriched for AcH3 (AcH3 Telo) +/− 4 kb. Graph below shows the sizes of peaks associated with AcH3 Telo sites were quantified. Significance was calculated using a two-way ANOVA, ns = not significant, *** = < 0.0005, **** = < 0.0001, ** = < 0.005, * < 0.05.

**Figure S8.**
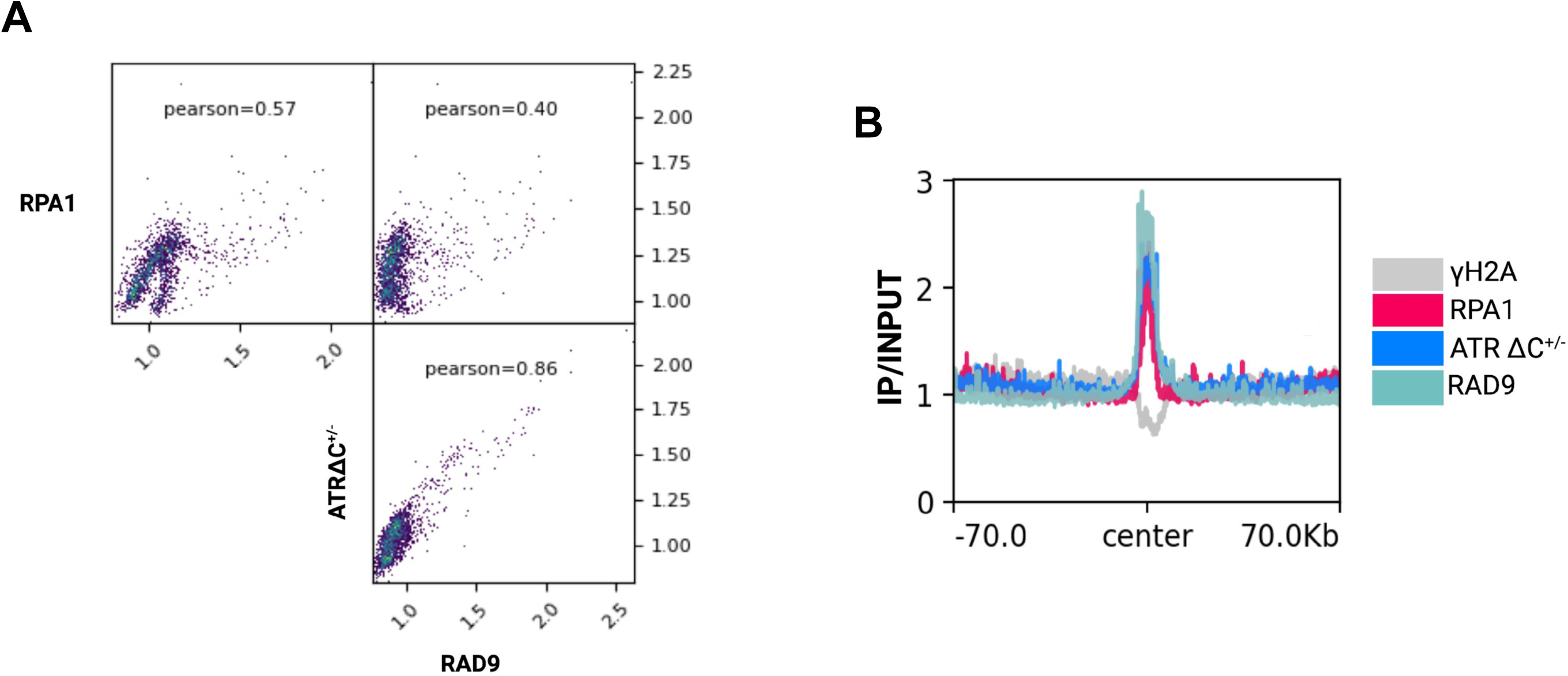
RPA1 and RAD9 localise to the early origin centre after exposure to 0.5 mM HU for 8 hrs. (A) Pearson coefficient correlation analysis to assess the global comparison of RPA1 ChIP-seq signal between RPA1 signal in RPAmyc and ATRΔC^+/−^ cells and RAD9myc ChIP-seq signal uncovered a poor correlation between RPA1 signal in ATR-competent and ATR-deficient cells. (B) Metaplot analysis showing the enriched signals for yH2A, RPA1 (in control and ATR-deficient cells) and RAD9 centred on the early origin site +/− 70,000 bp. ChIP-seq samples were normalised to their corresponding input controls.

**Figure S9.**
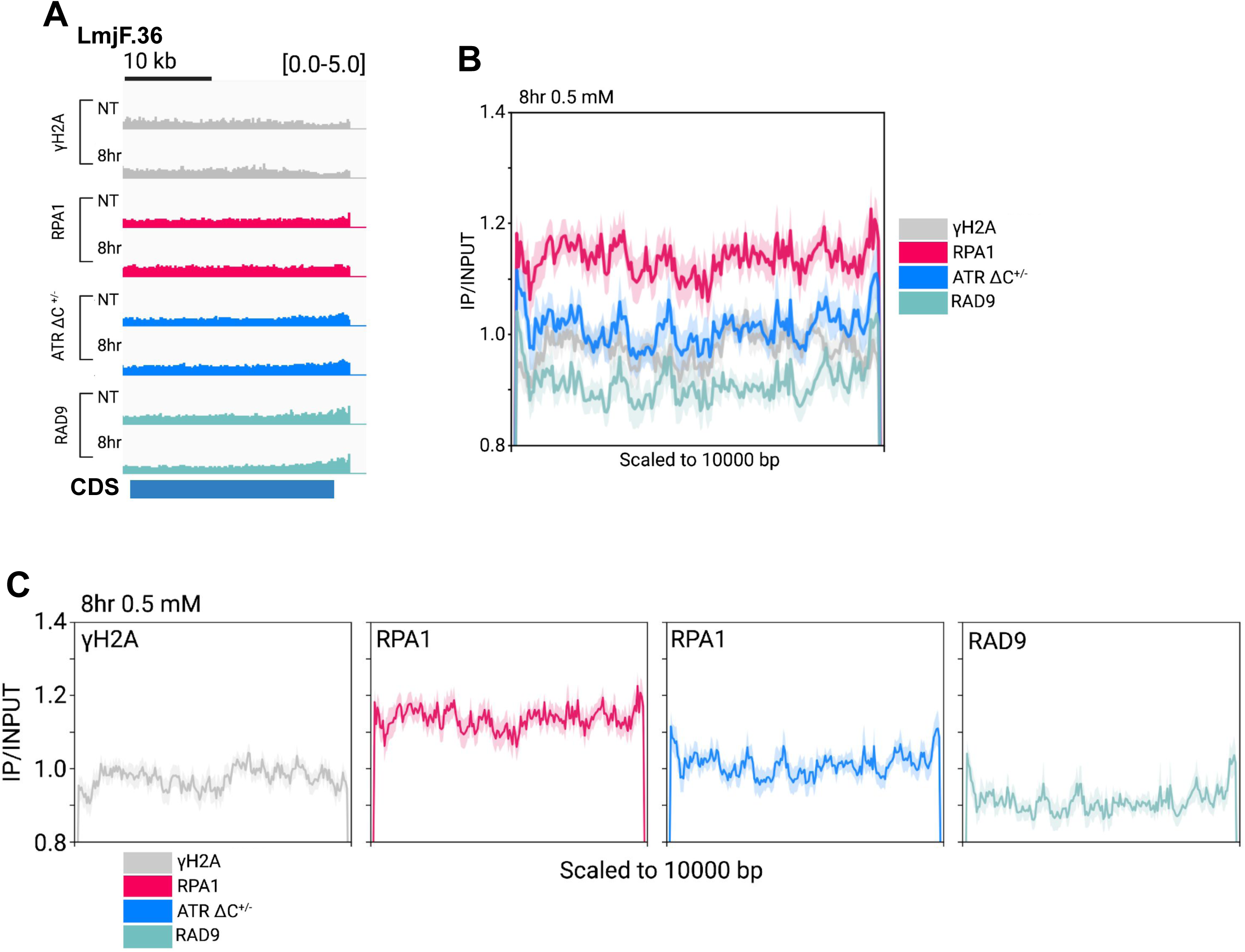
Subtelomeric accumulation of RS-associated factors in cells treated with 0.5 mM HU for 8 hrs. (A) Localisation of yH2A, RPA1 and RAD9 in control cells and the location of RPA1 in ATR-deficient cells. ChIP-seq signals shown are mapped to a representative subtelomeric region located on chromosome LmjF. 36. ChIP-seq samples were normalised to input. Scale = 0.0 – 5.0. Tracks were viewed in IGV. Polycistronic unit negative (blue). (B) Metaplot analyses showing the enriched signals for yH2A, RPA1 and RAD9 scale to subtelomeric sites (10,000 bp). ChIP-seq samples were normalised to their corresponding input controls. For clarity, individual metaplots for each protein are shown in (C). Signal is scaled as described in B.

